# *De novo* evolution of macroscopic multicellularity

**DOI:** 10.1101/2021.08.03.454982

**Authors:** G. Ozan Bozdag, Seyed Alireza Zamani-Dahaj, Thomas C. Day, Penelope C. Kahn, Anthony J. Burnetti, Dung T. Lac, Kai Tong, Peter L. Conlin, Aishwarya H. Balwani, Eva L. Dyer, Peter J. Yunker, William C. Ratcliff

## Abstract

While early multicellular lineages necessarily started out as relatively simple groups of cells, little is known about how they became Darwinian entities capable of open-ended multicellular adaptation^1,2^. To explore this, we initiated the Multicellularity Long Term Evolution Experiment (MuLTEE), selecting for larger group size in the snowflake yeast (*Saccharomyces cerevisiae*) model system. Given the historical importance of oxygen limitation^3^, our ongoing experiment consists of three metabolic treatments^4^: anaerobic, obligately aerobic, and mixotrophic yeast. After 600 rounds of selection, snowflake yeast in the anaerobic treatment evolved to be macroscopic, becoming ~2·10^4^ times larger (~mm scale) and ~10^4^-fold more biophysically tough, while retaining a clonal multicellular life cycle. They accomplished this through sustained biophysical adaptation, evolving increasingly elongate cells that initially reduced the strain of cellular packing, then facilitated branch entanglements that enabled groups of cells to stay together even after many cellular bonds fracture. In contrast, snowflake yeast competing for low oxygen remained microscopic, evolving to be just ~6-fold larger, underscoring the critical role of oxygen levels in the evolution of multicellular size. Taken together, this work provides unique insight into an ongoing evolutionary transition in individuality, showing how simple groups of cells overcome fundamental biophysical limitations via gradual, yet sustained, multicellular adaptation.

## Introduction

Organismal size plays a fundamental role in the evolution of multicellularity. The evolution of larger size allows organisms to gain protection from the external environment^5^ and explore novel niches^6^, while creating opportunities for the evolution of cellular differentiation^7–11^. Increases in organismal size have also been hypothesized to play a key role in the evolution of trade-off breaking multicellular innovations, as large size creates an evolutionary incentive to solve challenges of nutrient and oxygen transportation that are otherwise inescapable consequences of diffusion limitations^12,13^. However, little is known about how nascent multicellular organisms, consisting of small groups of undifferentiated cells, evolve to form biomechanically tough, macroscopic multicellular bodies, and whether selection for size itself can drive sustained multicellular adaptation^3^.

The evolution of macroscopic size presents a fundamental challenge to nascent multicellular organisms, requiring the evolution of biophysical solutions to evolutionarily-novel stresses that act over previously-unseen multicellular size scales^14–18^. While prior work with yeast and algae have shown that novel multicellularity is relatively easy to evolve *in vitro*, these organisms remain microscopic, typically growing to a maximum size of tens to hundreds of cells^19–23^. Extant macroscopic multicellular organisms have solved the above challenges through developmental innovation, evolving mechanisms that either reduce the accumulation of biophysical strain or increase multicellular toughness^24–26^. However, in nascent multicellular organisms that have not yet evolved coordinated morphogenesis, we do not know how, or even whether, simple groups of cells can evolve the increased biophysical toughness required for the evolution of macroscopic size.

Here we examine the interplay between biological, biophysical, and environmental drivers of macroscopic multicellularity using long-term experimental evolution. We subject snowflake yeast^21^, a model of undifferentiated multicellularity, to 600 rounds (~3,000 generations) of daily selection for increased size. Furthermore, because oxygen is thought to have played a key role in the evolution of macroscopic multicellularity, we evolved snowflake yeast with either anaerobic, mixotrophic, or obligately aerobic metabolism. All five of our replicate anaerobic populations evolved macroscopic size, while all aerobic and mixotrophic populations remained microscopic throughout the experiment, supporting the hypothesis that growth under low concentrations of oxygen constraint the evolution of large multicellular size^4^. Macroscopic size convergently evolved through two key changes in all five replicate populations. First, snowflake yeast increased the length of their constituent cells, which delays organismal fracture caused by packing-induced strain^18^. Next, they evolved to entangle branches of connected cells such that breaking a single cell-cell bond no longer causes multicellular fracture, evolving to become ~2·10^5^ times larger, forming millimeter-scale groups of clonal cells. Together these adaptations increased the toughness of individual clusters by more than 10^4^-fold, transforming the initial snowflake yeast ancestor, which was weaker than gelatin, to an organism with the strength and toughness of wood. Fitness assays, sequencing, and synthetic strain constructions reveal that macroscopic multicellularity evolved via selection acting on group size, an emergent multicellular trait of mutations directly affecting cellular morphology.

## Results

In 2018 we initiated the Multicellularity Long Term Evolution Experiment (MuLTEE), named after the pioneering Long Term Evolution Experiment with *E. coli* initiated by Rich Lenski^27^. This central goal of this project, which we intend to run over decadal time scales, is to observe open-ended multicellular adaptation in a nascent multicellular organism. We began the MuLTEE by engineering a unicellular isolate of *S. cerevisiae* strain Y55 to grow with the snowflake phenotype by deleting the *ACE2* open reading frame, ensuring that each replicate population had the same initial mechanism of group formation^28^. To examine the effect of oxygen on the evolution of size, we initiated three treatments in an otherwise isogenic ancestor: anaerobic growth (generated by selecting for a spontaneous petite mutant incapable of respiration), mixotrophy (cultured with glucose as the primary carbon source), and obligately aerobic growth (cultured with glycerol as the primary carbon source)^4^. We refer to the five replicate populations of anaerobic, mixotrophic, and obligately aerobic populations as PA, PM, and PO 1-5, respectively. We maintained strong directional selection favoring larger cluster size throughout the experiment by selecting for increasingly rapid sedimentation prior to transfer to fresh media (see Methods for details). We evolved these 15 populations over 600 rounds of growth and settling selection (~3,000 generations, Fig. 1A).

**Figure 1.**
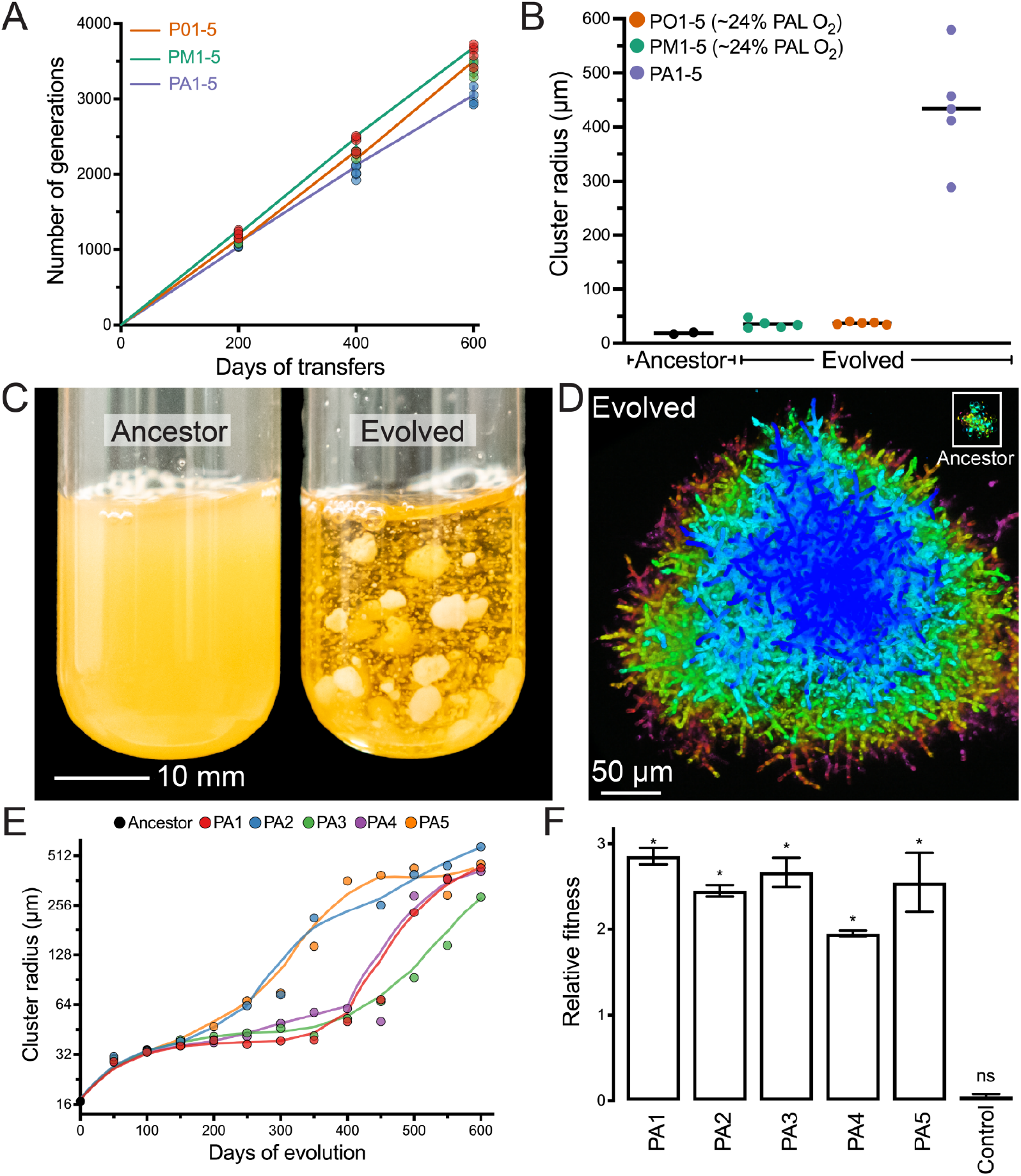
Evolution of macroscopic multicellularity in five replicate snowflake yeast populations. (A) We selected for larger size over 600 daily transfers, which represents ~3,000 generations. (B) Only the anaerobic populations (PA1-5) evolved macroscopic size over this time. (C) Individual snowflake yeast clusters from t600 are visible to the naked eye. (D) Representative clusters of the same two genotypes (ancestor in the upper right corner) shown under the same magnification (color represents depth in the *z* plane). (E) Temporal dynamics of size evolution in the anaerobic treatment (PA). (F) Macroscopic snowflake yeast were considerably more fit (calculated as a per-day selection rate constant (2.5)^30^) than their microscopic ancestor (*n*=3, error bars represent the SEM, asterisks denote significance at the 0.05 level). In A and E, the data points show the biomass-weighted mean radius (see Methods for details). See Extended Data Fig. 1 for additional data on the evolution of cluster size in oxygen-using populations (PM and PO) and Extended Data Fig. 3 for cluster size distributions for the 600-day anaerobic populations (PA1-5). Lines in (E) are Lowess smoothing curves intended to aid the eye.

All five populations of anaerobic snowflake yeast evolved macroscopic size, with individual clusters visible to the naked eye (Fig. 1B-D, Supplementary Movie 1). In contrast, snowflake yeast capable of metabolizing oxygen remained microscopic (Fig. 1A & Extended Data Fig. 1), a result consistent with recent work showing that competition for scarce oxygen imposes a powerful constraint on the evolution of large multicellular size^4^. Here, we focus on the evolution of macroscopic size in the five replicate anaerobic populations. Yeast in this treatment increased their mean cluster radius from 16 μm to 434 μm, a ~2·10^4^-fold increase in volume (Fig. 1E, *p* < 0.0001; *F*_5, 13321_ = 2100, Dunnett’s test in one-way ANOVA). This corresponds to an estimated increase from ~100 cells per cluster to ~450,000 (comparing average cluster volumes, accounting for changes in mean cell volume and cellular packing density within clusters).

The largest clusters of 600-day evolved macroscopic snowflake yeast are over a millimeter in diameter (Fig. 1C), which is comparable to the size of an adult *Drosophila^29^*. Much like their microscopic snowflake yeast ancestor^18,28^, macroscopic snowflake yeast possess a life cycle in which groups of cells both grow in size and reproduce, generating multicellular propagules, over the course of the ~24 h culture period (Extended Data Fig. 2). This analysis establishes that group size is heritable, and our time series data (Fig. 1E) show every replicate population evolved to form larger clusters at each 200-day sampling interval, strongly suggesting that larger size is an adaptive trait evolving in response to settling selection. To test this hypothesis, we performed a fitness assay competing the microscopic ancestor against each t600 PA1-5 population, under our standard selective conditions of growth and settling selection. The 600-day evolved macroscopic snowflake yeast were far more fit than their ancestor (mean daily selection rate constant = 2.5, Fig. 1F), increasing from a mean starting frequency of 52% to 99.9% over just three days.

As with their ancestor, macroscopic snowflake yeast grow via incomplete mother daughter cellular separation, forming a branched, tree-like structure (Fig. 2A&B). When compressed, macroscopic clusters fracture into small modules that resemble microscopic snowflake yeast in terms of branching morphology (Fig. 2 A&B). Their cellular morphology, however, changed markedly. Throughout the experiment, snowflake yeast cells evolve to be more elongate across all five replicate populations, increasing in average aspect ratio (ratio of length to width) from ~1.2 to ~2.7 (Fig. 2C&D; *F* 5, 1993 = 206.2, *p* < 0.0001, Dunnett’s test after one-way ANOVA; Extended Data Fig. 4). Even in macroscopic snowflake clusters, cell size and shape did not depend on location in the cluster (*i.e.*, interior or exterior; Extended Data Fig. 5). Initially, cluster size was a roughly linear function of cellular aspect ratio (Fig. 2E inset), but this relationship changes once they evolve macroscopic size (Fig. 2E).

**Figure 2.**
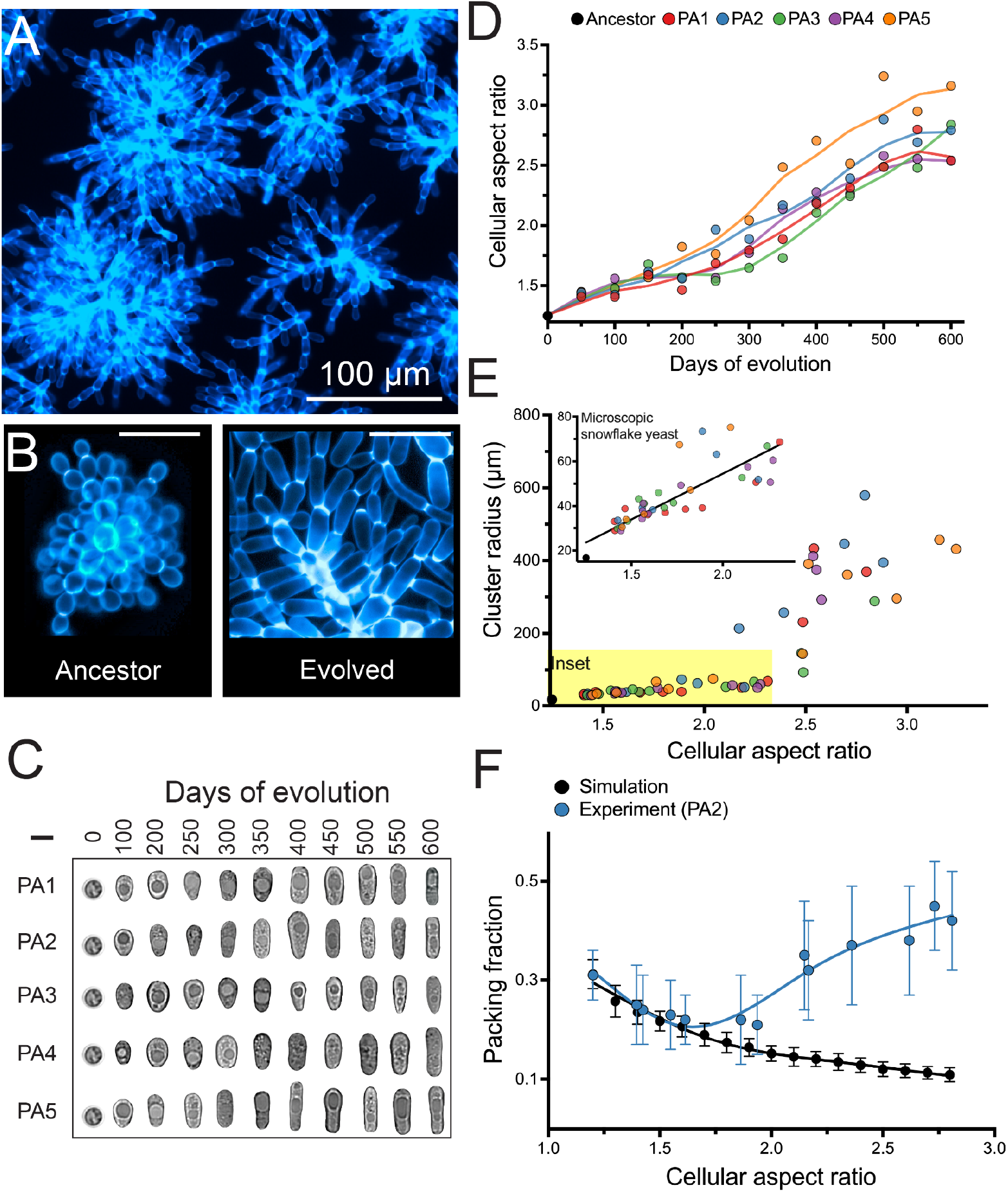
Evolution of novel cell morphology. (A) When compressed, macroscopic snowflake yeast fracture into modules, retaining the same underlying branched growth form of their microscopic ancestor as seen in (B) (scale bars are 20 μm). Cell walls are stained with calcofluor-white in (A) and (B). (C) and (D) show the parallel evolution of elongated cell shape, resulting in an increase in average aspect ratio from ~1.2 to ~2.7 (for each point in D, 453 cells were measured on average. The t0 ancestor is the same for PA1-5. Scale bar in C is 5 μm). An expanded version of (C) is shown in Extended Data Fig. 6. (E) Early in their evolution (aspect ratio 1-2.3), cluster size (weighted mean radius) is an approximately linear function of cellular aspect ratio (inset; *p* < 0.0001, *y* = 41.1*x* - 27.8, *r*^2^ = 0.72). This relationship does not hold beyond aspect ratio ~2.5. (F) A biophysical model of snowflake yeast predicts that increasing cellular aspect ratio should decrease cellular packing fraction (black points). We see a close correspondence with these predictions for low aspect ratios, but our experimental data diverges from model predictions for aspect ratios beyond 2. Each datapoint in (F) reports the mean of 15 snowflake yeast clusters or 25 replicate simulations, ± one standard deviation.

Prior work has shown that the evolution of more elongate cells increases the size to which microscopic snowflake yeast grow by decreasing the density of cellular packing (*i.e*., their packing fraction) in the cluster interior, which reduces cell-cell collisions that drive multicellular fracture^18,31^. To establish a null expectation for the effect of cell aspect ratio on cluster packing fraction, we simulated the growth of individual clusters from a single cell using an experimentally-validated model (see methods for details)^18^. In these simple 3D simulations, cellular packing fraction decreased monotonically with increasing cellular aspect ratio (Fig. 2F). We then examined this relationship over the course of our long-term experiment in replicate population two (PA2), which was one of the first lineages to evolve macroscopic size. As predicted by our simulation, cellular elongation decreased the packing fraction of microscopic multicellular groups— but only initially, from aspect ratio ~1.2-2.0. Beyond this, clusters with more elongate cells actually became more densely packed, and experimentally-measured packing fraction became increasingly divergent from model predictions (Fig. 2F). This divergence suggests that this lineage evolved a novel biophysical mechanism for increased multicellular toughness, capable of withstanding growth to macroscopic size and a high cellular packing fraction.

The simplest way that snowflake yeast could evolve to become macroscopic is to become adhesive, forming large aggregates of many separate snowflake yeast clusters. Indeed, aggregation is a common mechanism of group formation in yeast (*i.e*., via flocculation^32^), and this would explain the modular structure of macroscopic snowflake yeast (Fig. 2A). To determine if clusters of macroscopic yeast form via aggregation, or if they develop as a single clonal lineage, we labeled a single-strain isolate (taken from PA2 after 600 days of selection), with either GFP or RFP. If adhesive aggregation were responsible for their large size, we would expect to see chimeric groups composed of both red and green-fluorescent sub-clusters^32^. After five rounds of co-culture, however, all multicellular clusters (n=70; Extended Data Fig. 7) remained monoclonal. This is unlikely to occur with aggregation. If we conservatively assume each macroscopic snowflake yeast cluster we measured was the result of just a single fusion event, occurring with equal probability between two groups of red and green cells, then the binomial probability of finding no chimeric groups in our sample would be 10^-6^. Floc-like aggregation thus does not explain the evolution of macroscopic size in snowflake yeast.

To examine how changes in the topology of macroscopic snowflake yeast may underlie their increased size, we imaged clusters via Serial Block Face Scanning Electron Microscopy (SBF-SEM). This technique enables us to image the interior of macroscopic clusters that are difficult to resolve with light-based microscopy, allowing us to map their internal architecture with nanometer precision^33^. Within macroscopic clusters, separate branches contact, intercalate, and even wrap around each other (Fig. 3A). As these clusters are densely packed, moving one component would require moving many other components as well. Further, we found that individual macroscopic snowflake yeast were not composed of a single topologically-connected component, like their ancestors. Instead, they contained disconnected branches of cells, suggesting that the cluster remained intact even when cell-cell connections were broken (Fig. 3A). Based on these observations, we hypothesized that branches of macroscopic clusters are entangled, in a manner reminiscent of physical gels^34^ and entangled granular materials^35^. Entanglement would provide a mechanism for branches of cells to remain in the same, densely packed group even after cell-cell bonds break.

**Figure 3.**
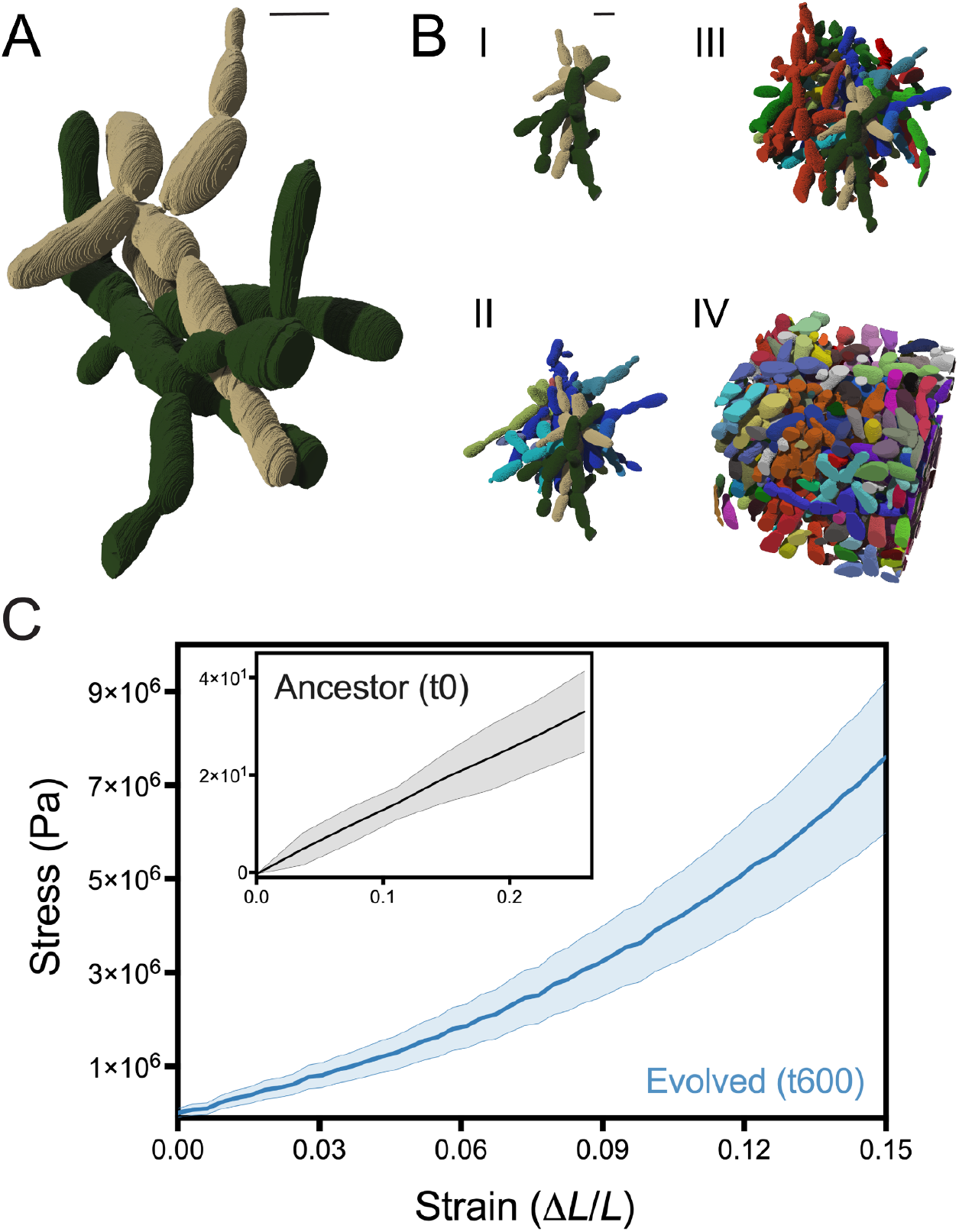
Branch entanglement underlies the evolution of macroscopic size. (A) Shown are two entangled components (green and tan), obtained via SBF-SEM imaging. (B) Branch entanglement is pervasive in macroscopic snowflake yeast. Starting with the two-component sub-volume in (A), we percolated entanglement by adding on adjacent entangled components in four steps (I-IV). Scale bars on A and B are 5 μm. (C) Stress vs. strain plot for macroscopic snowflake yeast (PA2, t600) clusters in blue and the ancestor in grey (ancestor shown again in inset with a rescaled *y* axis). Macroscopic snowflake yeast experience strain stiffening, a hallmark of entangled systems, while the ancestor’s stress-strain plot is linear, which is expected for non-entangled systems. The shaded area shows one standard deviation based on 10 repeated measurements for each.

Following prior work in entangled chains and knotted strings^35,36^, we used our SBF-SEM dataset to quantify branch entanglement in macroscopic snowflake yeast by analyzing chain topology and geometry. Specifically, we constructed the convex hull of each connected component within a sub-volume, which denotes the smallest convex polyhedron containing this component (see Extended Data Fig. 8A). If a cell from one connected component overlaps with the convex hull of a second, then the two can be considered entangled. By percolating entanglement among adjacent connected components throughout the sub-volume, we can measure the extent to which the cluster’s biomass is mutually entangled (Fig. 3B, Extended Data Fig. 8B). For entanglement to underlie macroscopic size, the largest entangled component (consisting of many entangled pieces) must be able to resist mechanical stress, meaning that there must be an entangled component that spans the vast majority of the cluster^37^. In analyses of 10 randomly selected sub-volumes from different macroscopic snowflake yeast clusters from population PA2 t600 macroscopic yeast, we found that the largest entangled component contains 93% +/- 2% of all connected components. This observation supports the hypothesis that entanglement between cell branches can prevent cluster fracture in the event that a cell-cell bond fractures.

As a further test, we investigated the mechanics of macroscopic snowflake yeast. Entangled materials are known to exhibit two key mechanical signatures: strain stiffening and high material toughness^35,38^. Strain stiffening describes the fact that, when compressed, the effective stiffness of entangled chains increases with increased strain. By efficiently distributing stress across constituent bonds, entangled materials can withstand stress orders-of-magnitude greater than their non-entangled counterparts^38,39^. As the microscopic ancestor is presumably not entangled, it should not exhibit strain-stiffening behavior or possess high toughness. We measured the mechanical stress response of 10 macroscopic snowflake yeast clusters under uniaxial compression using a macroscopic mechanical tester (Zwick Roell Universal Testing Machine). We repeated the same experiment for 10 ancestral microscopic snowflake yeast clusters using an atomic force microscope (AFM Workshop LS-AFM). The stress-strain plot for the microscopic ancestor is linear (r^2^ = 0.97 +/- 0.02, average and standard deviation of the regression for 10 samples, Fig. 3C inset), clusters fracture at stress as low as 240 Pa and have toughness as low as 8.9 J/m^3 18^. By contrast, macroscopic snowflake yeast clusters have a convex stress-strain curve (Fig. 3C), can support stresses at least as large as ~7 MPa without failing, and have toughness greater than 0.6 MJ/m^3^. Thus, entanglement both enables separate branches within macroscopic snowflake yeast to stay together and allows them to endure the large stresses necessary for growth to macroscopic size.

To rule out alternative hypotheses, we made additional measurements on macroscopic snowflake yeast (PA2 t600) and their microscopic ancestor. First, we measured the stiffness of individual cells to determine if they were becoming tougher. We did not detect a change between than ancestor and t600 macroscopic strain (0.018 and 0.019 N/m for the ancestor and evolved, respectively, Extended Data Fig. 9a). To determine if, in the absence of macroscopic entangled structures, individual PA2 t600 cells still show strain stiffening behavior, we crushed macroscopic clusters into smaller, microscopic branches before compressing them. These small groups displayed a linear stress-strain curve like their unevolved microscopic ancestor. In the absence of their macroscopic phenotype, the cells of the PA2 t600 yeast do not behave like an entangled material (Extended Data Fig. 9B).

Finally, we performed a positive control: we created persistently entangled groups experimentally. As described in Extended Data 7, growth in well-mixed liquid media prevents the formation of chimeric groups through entanglement. Yet if entanglement is critical for multicellular toughness, allowing fractured branches to remain in the same group, then chimeric clusters held together only by entanglement should be possible to grow under the right environmental conditions. We allowed GFP and RFP-tagged versions of PA2 t600 to grow at high density on solid media for 48 h, then cultured these yeast in liquid media for two rounds of growth and settling selection. These yeast readily formed and maintained chimeric groups. Specifically, 30% (31/101) of the clusters of the macroscopic genotype were still chimeric, with visibly entangled branches of green and red yeast (Extended Data Fig. 10). In contrast, only 1/110 clusters of the ancestral genotype were chimeric when cultured under the same conditions (0.9%, *p* < 0.001, *t* = 6.59, df = 209, two-tailed *t*-test). Taken together, this experiment shows that entanglement allows evolved snowflake yeast to remain intact, even when constituent branches lack continuous mother-daughter cellular bonds (*i.e.*, red and green branches are not attached to each other by permanent bonds).

To uncover the genomic basis of cell-level changes underlying multicellular adaptation, we sequenced the genomes of a single strain from each of the five populations (PA1-PA5) that independently evolved macroscopic multicellularity after 600 transfers (Fig. 4A&B). Over ~3,000 generations, snowflake yeast in our anaerobic treatment evolved dramatically more elongate cells (Fig. 2C&D), which plays a central role in the evolution of increased cluster size (Fig. 2E) and biophysical toughness (Figs. 2F & 3). Gene Ontology (GO) terms associated with cell length were significantly enriched, namely genes of the cell cycle^40^ (29 / 123 mutations, *p* = 0.02) and filamentous growth (7 / 123 mutations). In addition, we found 11 nonsynonymous mutations in genes with known roles in cellular budding (Fig. 4D), which includes eight genes that have previously been shown to increase the size of the bud neck (*AKR1, ARP5, CLB2, GIN4, PRO2, RPA49, RSC2, PHO81*)^41,42^. Mutations arose in two of these genes in different populations (*i.e*., *PHO81* in populations PA1 and PA5, and *GIN4* in populations PA2 and PA3, Fig. 4C), indicating parallel evolution. Larger bud scars should increase the amount of cell wall connecting cells, increasing the strength of the bond and toughness of the group. In our ancestral strain Y55, *gin4Δ* cells formed bud necks that had a 1.7-fold larger cross-sectional area (*t*=2.8, df = 8, *p* < 0.01). Consistent with this, we found that PA2-t600 macroscopic snowflake yeast evolved to form bud necks that had a 2.4-fold larger cross-sectional area (*t* = 5.3, df = 24, *p* < 0.001), and bud scars that were 5.8x larger 3D volume (*t* = 7.3, df = 24, *p* < 0.001) than their microscopic ancestor (Extended Data Fig. 11).

**Figure 4.**
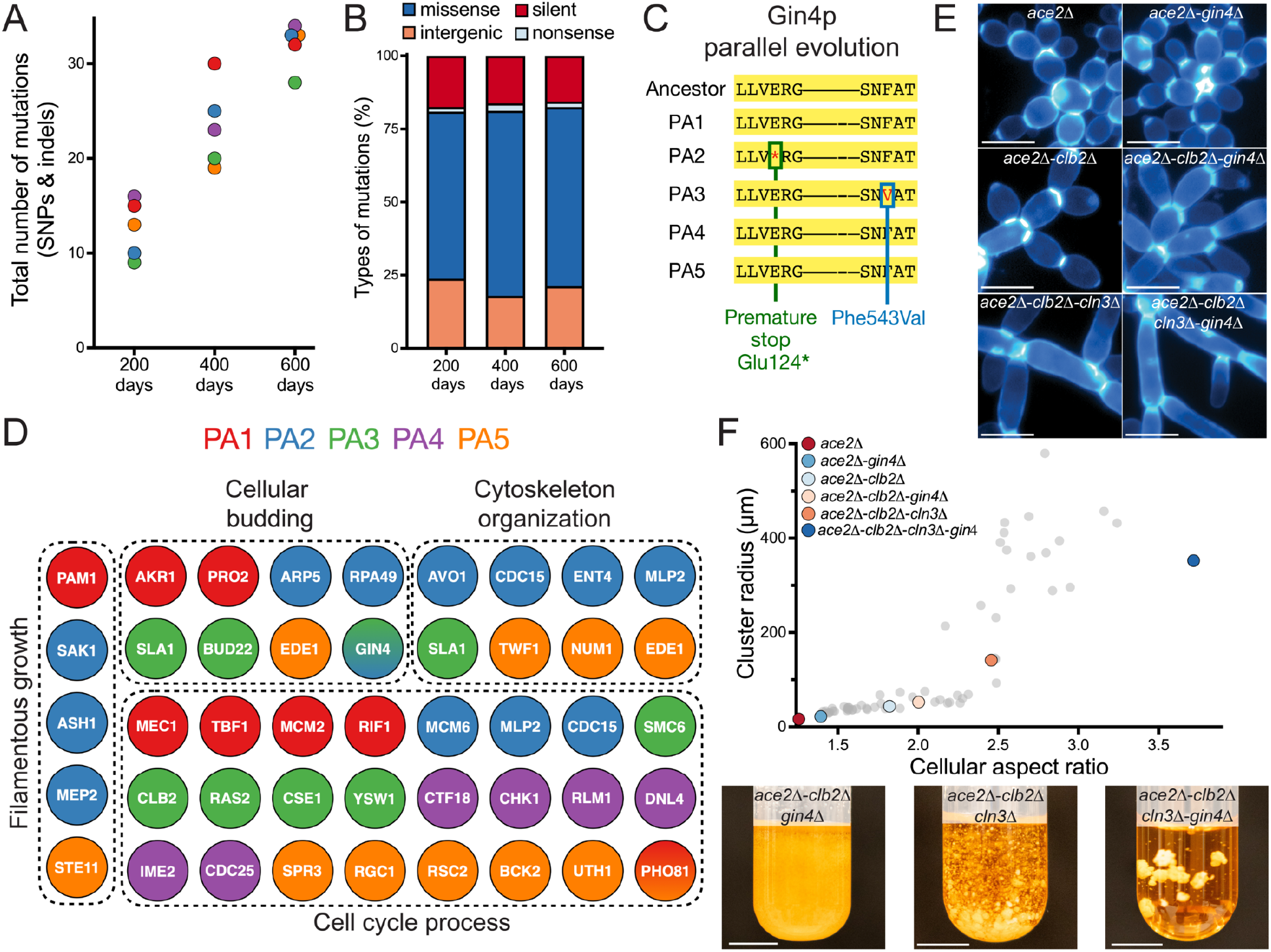
Whole-genome sequencing reveals the dynamics of molecular evolution and the genetic basis of cell-level and cluster-level changes. (A) and (B) show the number and types of mutations in evolved single strains from each population. (C) *GIN4*, a kinase whose loss of function increases bud neck size, is mutated in two independent populations. (D) Macroscopic snowflake yeast were enriched in mutations affecting cell cycle progression, cytoskeleton, and filamentous growth. In addition, we saw mutations affecting budding (*i.e.*, the location of buds on the cell surface, and bud neck size). These constituted 11 non-synonymous mutations out of the 123 total mutations in genotypes isolated from PA1-PA5 after 600 days of evolution. (E) Representative images of cells from strains used to re-engineer macroscopic size. Scale bars are 10 μm. (F) Engineered strains recapitulated the evolutionary trajectory established over 600 rounds of selection. With cellular aspect ratio below ~2.5, snowflake yeast remained microscopic, greatly increasing in size beyond this threshold. Scale bars are 10 mm.

As a final proof of principle, we set out to show that cellular elongation, aided by increased cell-cell bond strength, is sufficient to underpin the origin of macroscopic size in snowflake yeast.

Starting with the microscopic *ace2*Δ ancestor, we deleted the cyclins *CLN3* and *CLB2* in order to artificially increase cellular aspect ratio, and deleted *GIN4* to increase bud scar size, and thus strength. *CLN3*, while not present in evolved isolates, has a large, well-understood phenotypic effect on cell shape. Deleting *CLN3* and *CLB2* increased cellular aspect ratio (AR) by 21% and 45% respectively in single mutants, and 95% in the double mutant, with *GIN4* deletion further increasing the aspect ratio of each genotype in addition to its effects on bud scars (Fig. 4E and F). Our results mirror those from our evolution experiment: strains with AR < 2.5 were clearly microscopic, with increasing AR resulting in a gradual increase in group size. At AR ~2.5, *ace2Δ clb2Δ cln3Δ* yeast were at the threshold of macroscopic size, but still quite a bit smaller than our t600 isolates. The *ace2Δ clb2Δ cln3Δ gin4Δ* mutant, with an AR of 3.7, formed well-developed macroscopic clusters (Fig. 4F; experimentally-evolved strains described in Fig. 1E are shown in gray to facilitate direct comparison).

## Discussion

In this paper, we show that snowflake yeast, a model system of undifferentiated multicellularity, were capable of sustained multicellular adaptation, evolving macroscopic size over 600 days of experimental evolution. Macroscopic snowflake yeast are readily visible to the naked eye, containing hundreds of thousands of clonal cells. They achieved this remarkable increase in size by evolving highly elongate cells that become entangled within the cluster interior. This critical innovation allows multicellular groups to remain physically attached even when individual cellular connections are severed, increasing cluster toughness by more than 10,000-fold. As a material, snowflake yeast evolve from being ~100-fold weaker than gelatin^43^, to having the strength and toughness of wood^44^. Sequencing revealed an enrichment in mutations affecting the cell cycle and budding - traits that increase cell length and the amount of cell wall material at the point of attachment. Engineered strains with mutations increasing cell length and bud scar size recapitulated our evolutionary progression from microscopic to macroscopic size.

In our system, novel multicellular traits arise as an emergent property of changes in cell-level traits. Two cell-level innovations appear to have played a key role in the evolution of macroscopic size: more elongate cells and larger bud scars. Increased cell length initially reduces the strain generated from cellular packing, which is the primary manner in which size increased early in the experiment, and may underlie entanglement by facilitating cellular intercalation. Larger bud scars increase the amount of shared cell wall connecting cells, which all else equal should increase multicellular toughness by strengthening cell-cell bonds^31^. While the evolution of larger, tougher multicellular groups necessarily has underlying cell-level causation, these group and cell-level traits are distinct and non-commensurable (*i.e*., group size and toughness cannot be measured at the single-cell level). This demonstrates that snowflake yeast are evolving under MLS2, a shift in evolutionary dynamics that is critical for the transition multicellular individuality, as it allows groups as a whole, not just their constituent members, to gain adaptations^45,46^.

Entanglement is a common mechanism through which filamentous materials can solidify. It can operate on nearly any length scale, ranging from nanoscale polymers^47^ and nanofibers^48^, to macroscopic staples^49^, and beaded chains^50^. Relatively little is known about the role of entanglement in the materials properties of macroscopic biological structures, though recent work has shown that California Blackworm collectives are entangled, and can vary their degree of entanglement to solidify and melt their groups in response to environmental change^51–53^. Macroscopic multicellularity has evolved repeatedly in fungi^54^, and while to our knowledge no prior work has formally examined whether the cells of fungal fruiting bodies and lichen thalli are physically entangled, they are generally composed of densely-packed, overlapping hyphae, strongly suggesting entanglement^55–57^. The prevalence of entanglement in superficially different systems is likely due to its simplicity and efficacy; if pairs of constituents are easily entangled, large mutually-entangled clusters readily form, greatly increasing the strength and toughness of the material. While further work will be required to test this hypothesis, the relative ease with which multicellular fungi form entangled structures may have facilitated the highly convergent evolution of macroscopic multicellularity within this clade^54^, allowing different fungal lineages to independently evolve robust multicellular structures.

Our results depend on the fact that snowflake yeast grow as topologically-structured groups with permanent cellular bonds, and we would not necessarily expect similar biophysical exaptation in organisms with alternative means of group formation. These features, however, make it well suited as a model system for the lineages that have ultimately evolved complex multicellularity. Of the five lineages that independently evolved complex multicellularity (fungi, animals, plants, red algae, and brown algae), all but animals possess permanent cell-cell bonds, and early multicellular lineages in each are thought to have started out as simple, topologically-structured networks^58^. While animals do not currently have permanent cell-cell bonds, little is known about their ancestral mode of cellular adhesion. Indeed, their closest living relatives, the choanoflagellates, can form topologically structured multicellular groups with permanent cell-cell bonds^59–61^, suggesting that early animals may have possessed a similar mode of growth.

Despite 600 rounds of selection for increased size, our mixotrophic and obligately aerobic lineages remained microscopic (Fig. 1C & Extended Data Fig. 1), increasing their radius by less than two-fold. Extending prior work examining the role of oxygen diffusion in the evolution multicellular size^4^, these results highlight the importance of environmentally-dependent tradeoffs on the evolution of multicellularity. Oxygen can serve as a resource, allowing increased cellular growth by increasing ATP yields from metabolism^62^ and allows growth on non-fermentable carbon sources^63^. For simple, diffusion-limited organisms like snowflake yeast, low concentrations of oxygen create a cost to large size by reducing the proportion of cells in the group that have access to it- a cost which our anaerobic populations did not face.

During the evolutionary transition to multicellularity, groups of cells must become Darwinian entities capable of adaptation^64,65^. This requires that they reproduce, and have heritable variation in traits that affect fitness^66^. For groups of cells to become more than simply the sum of their parts, adaptation must take place in multicellular traits that are distinct from those of their constituent cells (*i.e*., they must evolve under MLS2)^45,46^. Until now, it has not been clear if simple groups of cells are capable of making this transition, or whether they first require innovations that allow for the heritable expression of novel multicellular traits^67,68^. Using long-term experimental evolution, we show even simple groups of cells, initially differing from their unicellular ancestor by a single mutation, have an innate capacity for open-ended multicellular adaptation. In response to selection on group size, a broadly important trait for simple multicellular organisms^69^, snowflake yeast evolved to form radically larger and tougher multicellular groups by leveraging the emergent biophysical properties of altered cellular morphology. These results demonstrate how selection on group size can drive sustained multicellular adaptation and biophysical innovation, and highlight the surprising ease with which evolutionary transitions in Darwinian individuality can occur.

## Methods

### Long-term evolution experiment

To generate our ancestral snowflake yeast for the MuLTEE, we started with a unicellular diploid yeast strain (Y55). In this yeast, we replaced both copies of the *ACE2* transcription factor using a *KANMX* resistance marker (*ace2::KANMX/ace2::KANMX*) and obtained a snowflake yeast clone (see^28^ for a detailed description of strains and growth conditions, including measurements of oxygen concentrations in growth media). When grown in YEPD media (1% yeast extract, 2% peptone, 2% dextrose), these yeast are mixotrophic, both fermenting and respiring. When grown in YEPG media, which is the same as YEPD but with the dextrose replaced by 2.5% glycerol, these yeast are incapable of fermentation and are obligately aerobic. From this initial clone of *αce2Δ* snowflake yeast, we selected a randomly produced ‘petite’ (p^-^) mutant. Due to a large deletion in its mitochondrial DNA (identified via sequencing), this snowflake yeast is unable to respire and is therefore metabolically ‘anaerobic,’ and was cultured in YEPD.

We evolved five replicate populations of mixotrophic snowflake yeast (referred to as populations PM1-PM5), obligately aerobic (PO1-PO5) and anaerobic (PA1-PA5) snowflake yeast in 10 mL of culture media, growing them in 25 x 150 mm culture tubes for 24 hours at 30°C with 225 rpm shaking. We used settling selection to select for larger cluster size. Once per day, after ~24 h of growth, we transferred 1.5 ml of culture into 1.5 mL Eppendorf tubes, let them settle on the bench for 3 minutes, discarded the top 1.45 mL of the culture, and only transferred the bottom 50 μl of the pellet into a new 10 mL of culture media for the next round of growth and settling selection. Once the anaerobic populations (PA1-PA5) had started to evolve visibly larger clusters with all biomass settling to the bottom of the tube in under a minute, we decreased the length of gravitational selection to 30 seconds, thus keeping them under directional selection for increased size. The timing of this change corresponded to ~350 days for PA2 and PA5 and ~500 days for PA1, PA3, and PA4. We used wide-bore filtered pipette tips (Thermo Scientific) for our daily transfers. In total, we applied 600 rounds (days) of growth and settling selection. We archived a frozen glycerol stock of each population at −80°C every 10-15 transfers.

### Measuring cluster size

We developed a standard visualization protocol to be able to measure the size of both microscopic and macroscopic snowflake yeast from each population over the 600-day evolution experiment. To prepare yeast for imaging, we revived evolved frozen cultures for each population in 50-day intervals (12 for each of the 15 replicate populations). We then inoculated each sample into 10 mL fresh media and brought them to equilibrium over a five-day culture process, performing daily settling selection prior to transfer to fresh media. After 5 transfers, we pipetted a random 1mL subsample of each 24-hour culture, placing them in 1.5 ml Eppendorf tubes. We added 0.5 ml of sterile water to each well of 12-well culture plates, then gently vortexed each snowflake yeast sample and diluted them into the water (1,000-fold dilution for microscopic populations, and 100-fold dilution for macroscopic populations). We shook each well plate gently to disperse the yeast clusters evenly over the bottom of each well. We then imaged each well using a 4X Nikon objective, capturing the cross-sectional area of clusters without disrupting their 3D structure. Next, we used ImageJ-Fiji to calculate the cross-sectional area of each cluster, converting pixels to microns by including a physical 100 μm scale bar in each image.

### Calculating weighted average cluster size

The distribution of cluster size across various isolates are not consistent-while microscopic populations are unimodal, while macroscopic populations contain a substantial number of small groups that may only contain a few cells. Even when these small groups constitute a trivial amount of the population’s biomass, variation in their abundance can have a large impact on sample statistics, like average cluster size. Because of their skewed size distribution, mean size is an unreliable and often uninformative measure of the central tendency of the cluster size distribution, and does not accurately describe how cells are distributed across different cluster size classes. To account for this, we calculated the distribution of cellular biomass over the range of cluster sizes, and found the mean of this biomass distribution (which is the same as weighting mean cluster size by its biomass). This weighted mean cluster size represents the expected size group any given cell will be in (see Extended Data Fig. 2 for a visual representation), and is an accurate measure of changes in the distribution of cellular biomass across different cluster sizes over evolutionary time. Rather than presenting the weighted mean group size as a volume, we transformed these into an average (micron) radius to be consistent with the units that have historically been used in the paleontological literature documenting the evolution of macroscopic multicellular organisms.

### Assessing fitness

We measured the relative fitness of the evolved macroscopic populations (PA1-5) in competition against the ancestor in liquid culture under the same conditions as our evolution experiment. To differentiate competing strains, we used an ancestral snowflake yeast strain carrying a hemizygous red fluorescent protein (*ura3::dTOMATO/URA3*). Before coculturing these strains, we first grew evolved populations and the ancestral strain in separate cultures overnight. Then we mixed the two types in a 1.5 mL microcentrifuge tube to start the competition assay in fresh 10 mL YEPD cultures. We examined the fitness of PA1-5 t600, as well as an ancestor:ancestor control, with three replicate competitions per treatment. We grew these competitions for 24 hours in 10 mL YEPD (conditions as described in the evolution experiment), followed by 3 minutes of settling selection in 1.5 mL Eppendorf tubes. We then transferred the bottom 50 μl into a fresh culture tube for the next round of growth and settling, repeating the same procedure for three rounds across the fitness assay. The evolved populations’ initial frequency ranged between 35-70%, and after three rounds of growth and settling selection, they reached a range between 99.8-99.9%.

To calculate the relative fitness of the evolved populations against the common ancestor, we calculated their selection rate constant, as described in ^30^. To do so, we estimated the initial and final cellular density of yeast by measuring the cross-sectional area of the evolved and ancestral snowflake yeast clusters at the beginning and end of the fitness assay using a Nikon Eclipse Ti inverted microscope at 100x magnification. Then we calculated the selection rate by dividing the estimated density of the evolved populations at the end and beginning of the competition assay, followed by subtracting the natural log of this value from that of the ancestral strain^30^. Finally, by coculturing the ancestral snowflake yeast strains with and without the hemizygous *dTOMATO* constructs, we confirmed that the expression of this protein had no significant fitness cost, as reported in Fig1F, right column (p=0.22, t=1.7, df=2, one sample t-test). Image analysis was performed in ImageJ (v2.3.0).

### Aspect ratio data collection and analysis

To measure the evolution of cellular aspect ratio in populations PA1-PA5 over the 600-day evolution experiment, we first inoculated 61 samples (1 ancestor + 5 replicates x 12 time points, each separated by 50 days) and grew them overnight in shaking incubation as described above. Following the same growth protocols as in our cluster size measurements, we grew these samples for five consecutive days with settling selection. On the final day, we transferred 100 μl of each culture into tubes with fresh YEPD and incubated them for 12 hours. Next, we stained samples in calcofluor-white by incubating them in the dark for 30 minutes (at a final concentration of 5 μM) prior to imaging (40x objective, UV excitation of blue fluorescent cell wall stain, imaged on a Nikon Ti-E). We measured the aspect ratio of individual cells within snowflake yeast clusters on ImageJ-Fiji, analyzing an average of 453 cells per population.

### Simple biophysical model examining packing fraction as a function of aspect ratio

We simulated the growth of snowflake yeast from a single cell. Cells were modeled as prolate ellipsoids with one long (major) axis and two equal shorter (minor) axes. Clusters started as a single cell and were grown for nine cellular generations. New cells first emerged from their mother’s distal pole; subsequent cells emerged with a polar angle of 45° and a random azimuthal angle. If adding a new cell would cause too much overlap with existing branches, the new cell was deleted and the mother cell lost its chance to reproduce that generation. We simulated the growth of 50 clusters of each genotype, which varied in their cellular aspect ratio, defined as the ratio of the major axis to minor axis length, ranging from 1.2-2.8. We then calculated each simulated cluster’s packing fraction by fitting a convex hull to the cluster, and measuring the ratio of the total volume to the volume specifically occupied by cells. The MATLAB code to grow snowflake yeast using this protocol is attached as Supplementary File 1.

### Testing aggregative vs. clonal development

To determine if macroscopic snowflake yeast aggregate or develop clonally (Extended Data Fig. 7), we isolated a single genotype from PA2, t600 (strain GOB1413-600), and engineered it to constitutively express either green or red fluorescent proteins. To do that, we amplified the *prTEF_GFP_NATMX* construct from a pFA6a-eGFP plasmid and the *prTEF_dTOMATO_NATMX* construct from a pFA6a-tdTomato plasmid. We then separately replaced the *URA3* open reading frame with *GFP* or *dTOMATO* constructs in an isogenic single strain isolate following the LiAc transformation protocol^70^. We selected transformants on Nourseothricin Sulfate (Gold Biotechnology Inc., U.S.) YEPD plates and confirmed green or red fluorescent protein activity of transformed macroscopic clusters by visualizing them under a Nikon Eclipse Ti inverted microscope. To test whether they grow clonally or aggregatively, we first inoculated *GFP* or *dTOMATO* expressing clones individually overnight. We then mixed the two cultures in equal volume and diluted 100-fold into a 10 mL fresh culture. We co-cultured this mixed population for five days, transferring 1% of the population to fresh media every 24 h. Finally, we washed this culture in 1 mL sterile water and visualized 70 individual clusters under both red and green fluorescent channels, allowing us to count the number of snowflake yeast clusters that were green, red, or chimeric.

We examined the potential for entanglement alone to allow for persistent interactions among disconnected components (Extended Data Fig. 10) by crushing GFP- and RFP-tagged macroscopic snowflake yeast (PA2-t600) into smaller groups, and then growing a mixture of them on the surface of agar plates for 48 hours, potentially allowing branches of adjacent genotypes to entangle through growth. We then scraped these populations and grew them in 10 mL YEPD (yeast-extract, peptone, dextrose) media with shaking at 250 rpm for two 24 h rounds of growth and settling selection. We imaged the resulting clusters under widefield microscopy (Nikon Ti-E), taking pictures of individual clusters under bright field, green, and red channels, collecting data for a random sample of clusters (101 for the PA2-t600, and 110 for the ancestor control). We analyzed the images in ImageJ-FIJI, and any clusters that contained both green and red cells were scored as chimeric. The images shown in Extended Data Fig. 10 were taken on a Nikon AR1 confocal, allowing better view of the 3D structure of chimeric intercalation.

### Specimen preparation for Serial Bulk Faced Scanning Electron Microscopy (SBF-SEM)

We fixed snowflake yeast in 2% formaldehyde (fresh from paraformaldehyde (EMS)) containing 2 mM CaCl2, incubating at 35°C for 5 minutes followed by 2-3 hours on ice. Next, we incubated these yeast for an hour in a solution of 1.5% potassium ferrocyanide, 0.15M cacodylate buffer, 2 mM CaCl2, and 2% aqueous osmium tetroxide. This last step was performed on ice and under vacuum. Finally, we washed our yeast and incubated them in thiocarbohydrazide solution (10 g / L double-distilled water) for 60 minutes at 60°C, followed by en bloc uranyl acetate and lead aspartate staining^71,72^.

### SBF-SEM

We imaged fixed yeast on a Zeiss Sigma VP 3View. This system has Gatan 3View SBF microtome installed inside a Gemini SEM column. For this work, yeast clusters that were embedded in resin were typically imaged at 2.5 keV, using 50-100 nm cutting intervals, 50 nm pixel size, beam dwell time of 0.5-1 μsec and a high vacuum chamber.

### SEM Image analysis

Images were initially in.dm3 format, which we converted to .tiff using GMS3 software. We then cleaned the images and passed them through a gaussian filter in Python. Using the interactive learning and segmentation toolkit (ilastik), we segmented images into 3 parts: live cells, dead cell debris, and background. We then imported segmented HDF5 files in Python. First, we identified connected cells using the nearest neighbor algorithm to identify connected cells. We call a set of connected cells inside a sub-volume a connected component. Then, using a 3D extension of the gift-wrapping algorithm, we extracted the convex hull of each connected component.

### Visualization of SEM images

After segmenting images as described above, we created a mesh of individual cells by dilating binarized images. After creating the surface mesh of each individual cell using the mesh tool in Mathematica 12, we imported whole sub-volumes in Rhino6. Then we manually identified cell-to-cell connections and colored each connected component differently.

### Volume fraction data collection and analysis

We measured the packing fraction (proportion of the cluster volume that is cellular biomass) by measuring the number of cells within a cluster, their size, and the volume of the cluster, following the protocol described in Zamani et al. (2021)^73^.

### Mechanical testing

To test the response of ancestral clusters to uniaxial compression we submerged individual clusters under water, and then compressed them using a Puima Chiaro nanoindenter (Optics11, 19.5 um spherical glass probe). For mechanical measurements of macroscopic snowflake yeast, we used a Zwick Roell Universal Testing Machine (UTM) with 5 N probe. As above, individual clusters were extracted from the growth tube and placed on the testing stage while submerged under water.

### Preparing glass slides with attached cells

We coated glass slides with Concanavalin A to make a sticky glass surface to which individual cells could adhere. We started by preparing a 10 mg/mL solution of ConA dissolved in sterile DI water, which can be stored at −20C. This stock solution was diluted 1:10, and then 200uL of diluted solution was pipetted onto a glass slide in a sterile environment. The slide was allowed to incubate for 5 min at room temperature, then washed with sterile DI water twice, then left to aspirate dry in the hood. Cell cultures were inoculated (100 uL) onto the glass surface and left to settle for 5 minutes.

### AFM measurements

Prior to measuring the properties of individual cells with the AFM, we restored ACE2 functionality to increase the frequency of single cells available for mechanical testing. To do this, we reinserted a single copy of the ancestral *ACE2* allele fused with the antibiotic resistance gene *HYGNT1* into the genome of the PA ancestor and PA1 t600 isolate under the control of its native promotor using the LiAc/SS-DNA/PEG method of yeast transformation^69^. Transformants were then plated on YEPD agar plates (1% yeast extract, 2% peptone, 2% dextrose, 1.5% agar) supplemented with 200 mg l^-1^ of the antibiotic hygromycin B (Enzo Life Sciences). All atomic force measurements used an atomic force microscope from Asylum Research that was integrated with an inverted optical microscope (Nikon). For single-cell measurements, we used a silicon nitride cantilever with a nominal stiffness of 0.06 N/m with an attached borosilicate glass bead with diameter 2um (Novascan Technologies). The cantilever was measured via thermal analysis to have a stiffness of 0.0593 N/m. For cluster-level measurements, we used tipless, aluminum coated cantilevers that have a rectangular shape (length 225 um, width 40 um) that have a nominal stiffness of 30 N/m (AppNano). For measurements, either single cells or entire clusters were visually aligned with the cantilever probe, which was then moved at a velocity of 1 um/s to compress the cell or cluster with increasing force.

### Chitin staining protocol

We stained cells with calcofluor white via the following protocol. First, we mixed 500uL of cell culture from the ancestor and PA2 t600 strains into the same tube. Then, we sampled 150 uL (containing both the ancestor and t600 yeast clusters) from the mixed culture. We removed the supernatant via an iterated process of centrifugation and pipetting media removal. Then, we diluted 15 uL of 1 mg/mL calcofluor solution into 500 uL 1x Phosphate-buffered saline solution (PBS) and mixed with the yeast pellet. We incubated the sample in darkness at room temperature for 25 minutes, then we removed the calcofluor media via centrifugation and pipetting. Finally, we added 200 uL 1x PBS on top of the pellet. 20 uL of this cell suspension was pipetted onto a clean glass slide and covered with a coverslip for microscopy.

### Single Cell and Bud Scar Confocal microscopy

We used a Nikon A1R confocal microscope equipped with a 60x oil immersion objective to obtain z-stack images of individual cells stained with calcofluor white. To track the location and size of bud scars, we wrote a MatLab script to extract the brightest calcofluor signals, since the chitinous bud site region makes bud scars brighter than the other portions of the cell wall. Brightness isosurfaces isolated the bud scars themselves, and the brightness of each voxel contained within the isosurface was recorded to track the density of chitin. Next, the isosurface points were rotated to the x-y plane by finding the principal axes of the shape via principal component analysis. The rotated surface points were then used to calculate the height and cross-sectional area of the bud scars.

### DNA extraction and genome sequencing

To extract DNA for whole-genome sequencing, we isolated clones from each of the evolved replicate populations of anaerobic yeast (*i.e*., PA1-PA5) and their common ancestor after 200, 400, and 600 days of evolution. To pick clonal isolates, we diluted populations of snowflake yeast clusters in 1.5 mL tubes and plated them at a density of 100-200 colonies per plate. Next, we restreaked those initial single colonies onto fresh plates, thus ensuring that each colony on a plate results from a single snowflake yeast cluster. Because snowflake yeast grow clonally, we expected that these isolates would only represent a single clone of cells, with no more variation than would be expected from any single cell isolate that grew into a population, generating *de novo* mutation along the way (subsequent analysis of the genomes confirmed this: we never saw evidence of >1 genotype present in any isolate). We inoculated these 16 samples in YEPD for 12 hours and extracted their genomic DNA using a commercially available kit (Amresco, Inc. VWR USA). We measured the concentration of DNA with a Qubit fluorometer (Thermo Fisher Scientific, Inc.). We prepared genomic DNA libraries for the 16 samples using NEBNext Ultra DNA Library Prep Kit for Illumina (New England Biolabs, Inc). We quantified the quality of the genomic DNA library using the Agilent 2100 Bioanalyzer system located at the Genome Analysis Core Laboratories at Georgia Institute of Technology (Agilent Technologies, Inc). Finally, whole genomes were sequenced using the HiSeq 2500 platform (Illumina, Inc) by the Genome Analysis Core Center located in the Petit Institute, Georgia Tech. As a result, we obtained paired 150 bp (R1 & R2) FASTQ reads from two lanes (L1 & L2).

### Bioinformatic analysis

For our bioinformatic analysis, we used the bash command-line interface on a Linux platform. To identify *de novo* mutations (single nucleotide changes, or ‘SNPs,’ and small insertion/deletions, or ‘indels’) in the ancestral and evolved genomes, we first filtered out low-quality reads using a sliding window approach on Trimmomatic (v0.39). We aligned reads to the yeast reference genome (S288C, SGD) using an algorithm in the BWA software package (*i.e.*, BWA-MEM)^74^. Next, we used the genome analysis toolkit (GATK) to obtain and manipulate .bam files^75^. Duplicate reads were marked using the Picard - Tools (MarkSuplicates v2.18.3). We called SNPs using two different tools, i.e., GATK4 HaplotypeCaller (v4.0.3.0) and FreeBayes (v1.2.0)^75,76^. We validated SNP calls by comparing results obtained by two independent tools. For indels, we used the output from HaplotypeCaller. To filter variants according to their quality/depth scores and generate an overview of the variant calling step’s statistical outcome, we used VCFTOOLS (v0.1.16)^77^. Finally, after manually checking each variant call by visualizing SAM files and VCF files on Integrative Genomics Viewer (IGV)^78^, we extracted *de novo* variants by making a pairwise comparison of each VCF file of the evolved genomes against the VCF file of the ancestral genome by using bcftools-isec (v1.10)^79^. Lastly, we annotated evolved mutations using SnpEff (v4.3T)^80^.

To search for gene ontology (GO) term enrichment for *de novo* mutations, we generated a combined list of synonymous and nonsynonymous mutations within gene coding regions. We then searched for enriched gene ontology terms using GO Term Finder and GO Slim Mapper on the yeast genome database^81^.

### Genetically engineering macroscopic snowflake yeast

To genetically engineer snowflake yeast strains with cell lengthening (*CLB2* and *CLN3*) and bud-scar strengthening (*GIN4*) mutations that are shown in Fig. 4, we used homozygous unicellular (GOB76) and multicellular strains (GOB8). To engineer each gene deletion, we first amplified hygromycin, nourseothricin, and G418 resistance marker cassettes from plasmids and individually transformed them into yeast cells (see Supplementary Table 1 for genotypes, plasmids, and primers). Next, we induced sporulation of individual heterozygous mutant strains (2% KAc), dissected tetrads, and obtained homozygous deletions through auto-diploidization. By subsequent rounds of sporulation and inter-tetrad mating on appropriate multi-drug plates, we generated double, triple, and quadruple mutant strains (Fig. 4 E&F). Finally, we quantified the cellular aspect ratio and cluster size (cross-sectional area) of these mutants by imaging clusters under a Nikon Eclipse Ti inverted microscope, using the same methods as previously described.

### Life-cycle experiment (Extended Data Fig. 2)

To characterize the life cycle of the ancestral (microscopic) and evolved (macroscopic, PA2-600) snowflake yeast, we inoculated both strains starting from frozen glycerol stocks. Then, we grew them under the conditions of the evolution experiment for five rounds of growth and settling selection. Applying a five-day growth and settling selection brings cultures to an equilibrium, reflecting the physiology and size distribution observed during the evolution experiment. On the final day, we sampled from the growing cultures at 0, 3, 6, 12, and 24 hours and measured the number, size, and volume of cultures using the same methods described under “Measuring cluster size.”

## Acknowledgements

We thank Jennifer T. Pentz for teaching us Illumina library preparation, Shweta Biliya at the High Throughput DNA Sequencing Core at Georgia Tech for sequencing the genomes of evolved strains, and Kingsley A. Boateng at Core Facilities at the Carl R. Woese Institute for Genomic Biology for the SEM imaging. Siyi Cao helped us with microscopy during the early stages of this project. We thank Christian Orlic, László Nagy, and all members of the Ratcliff group for insightful comments on the manuscript. This work was supported by NIH grants R35-GM138030-01 to W.C.R. and R35-GM138354-02 to P.J.Y., and a Packard Fellowship for Science and Engineering to W.C.R.

## Author Contributions

G.O.B., A.Z., P.J.Y, and W.C.R. conceived of the project. G.O.B. and W.C.R. designed the MuLTEE, and G.O.B. performed the evolution experiment. G.O.B., A.Z., P.K., and T.C.D. designed and collected data. A.Z. generated SBF-SEM images. A.Z., T.C.D., and P.J.Y performed the yeast biophysical simulations. E.L.D. and A.H.B. assisted G.O.B. and A.Z. with image analysis. A.J.B. genetically engineered large snowflake yeast, K.T. performed life-cycle experiments, D.T.L. measured the number of generations, and P.L.C. performed unicellular reversion experiments. G.O.B., A.Z., T.C.D., W.C.R., and P.Y. analyzed the data. G.O.B. made the figures. G.O.B., W.C.R., and P.J.Y. wrote the first draft of the paper, and all authors contributed to the revision.

## Data and code availability

All source data and code are available at https://github.com/ozanbozdag/de_novo_evolution_of_macroscopic_multicellularity. Microscopy images are available upon request.

## Extended Data Figures

**Extended Data Figure 1.**
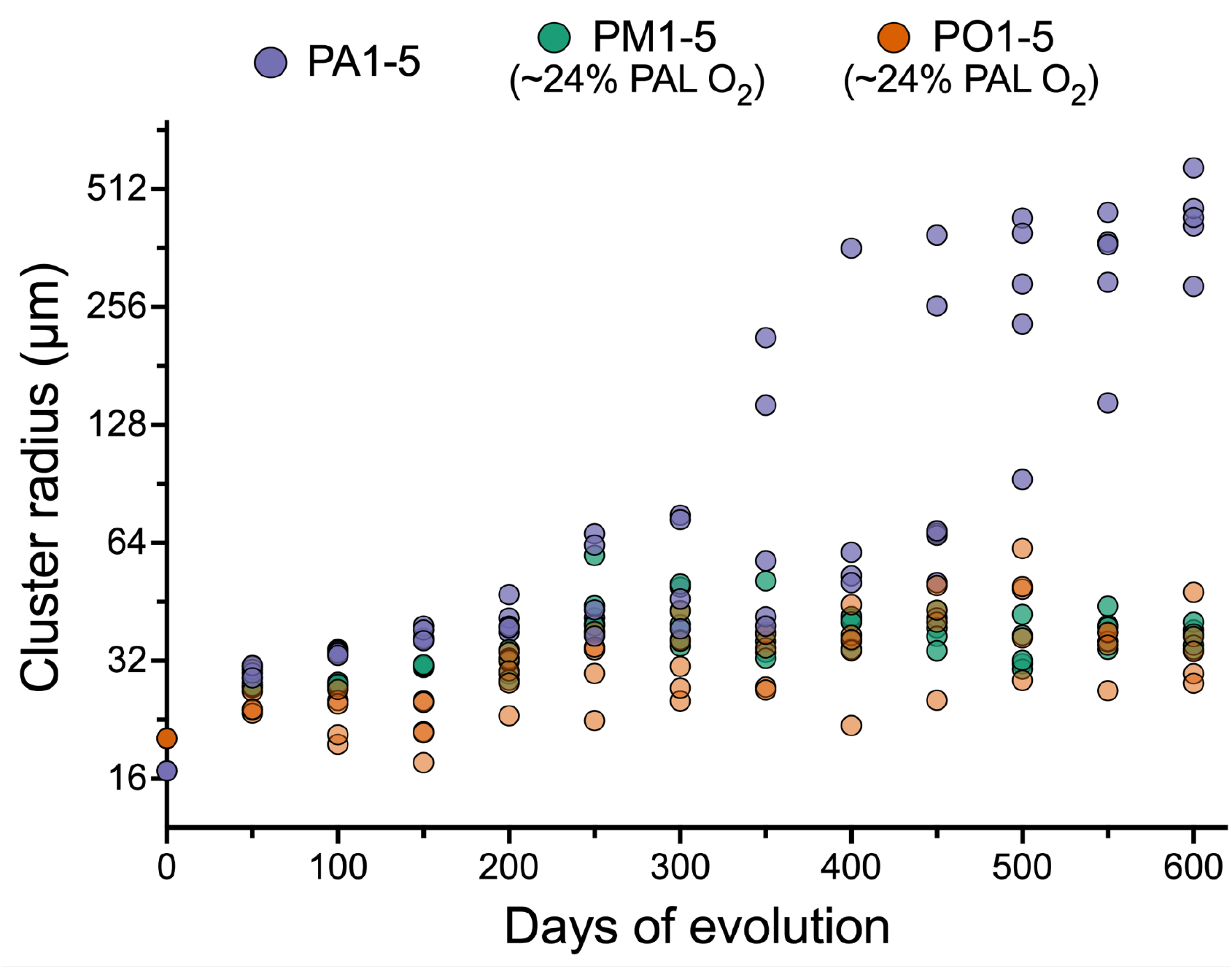
Temporal dynamics of size evolution in each population and treatment group. Data points show the weighted average radius of cluster size for the whole population (see Methods for details).

**Extended Data Figure 2.**
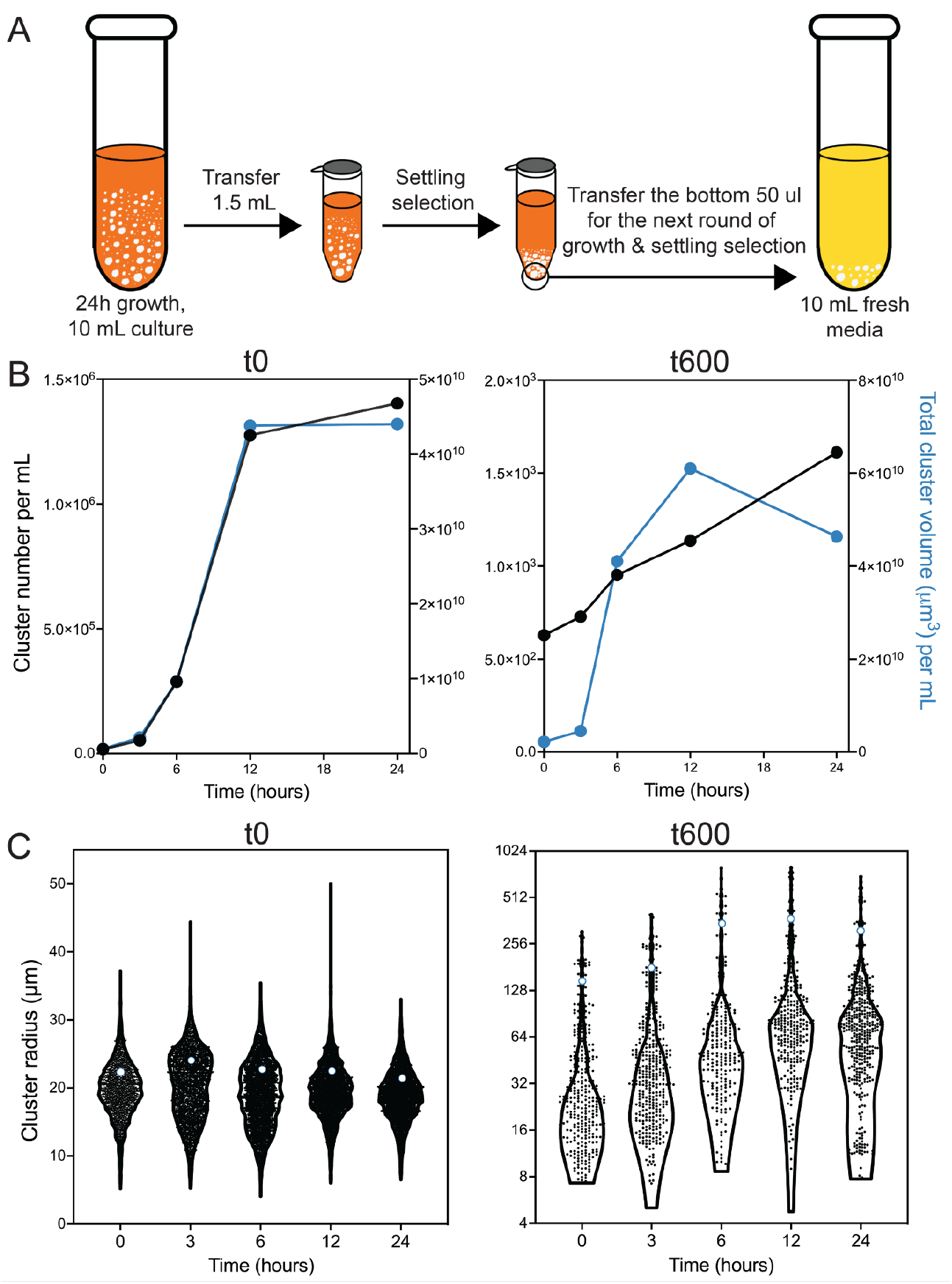
Characterizing the life-cycle of the ancestral (microscopic) and evolved (macroscopic) snowflake yeast. A) During the ~24-hour growth cycle, snowflake yeast compete for growth and reproduction in 10 mL of YPED (250 RPM at 30°C). At the end of the growth phase, we select for larger group size via settling selection. While there is a theoretical maximum survival rate of 15% (that is, if all of the cells survived settling selection), we only transfer the bottom 50 μl of pellet biomass regardless of how many cells settle, creating an arms race that favors the fastest groups within the population. Our measurements of the number of cellular generations per day in Figure 1A suggests about 3% of the cells survive from one day to the next on average. B) Both the microscopic (ancestral) and macroscopic (t600) snowflake yeast clusters have a life cycle, reproducing during the growth phase. C) Consistent with entanglement producing tough groups, macroscopic snowflake yeast release mostly microscopic propagules, possibly from branch tips at the exterior of the group, where the opportunity for entanglement is minimal. Despite the presence of many small propagules, most of the biomass in the population is contained within macroscopic clusters. The open circles represent the biomass-weighted mean size, which is the average sized group the mean cell finds itself in.

**Extended Data Figure 3.**
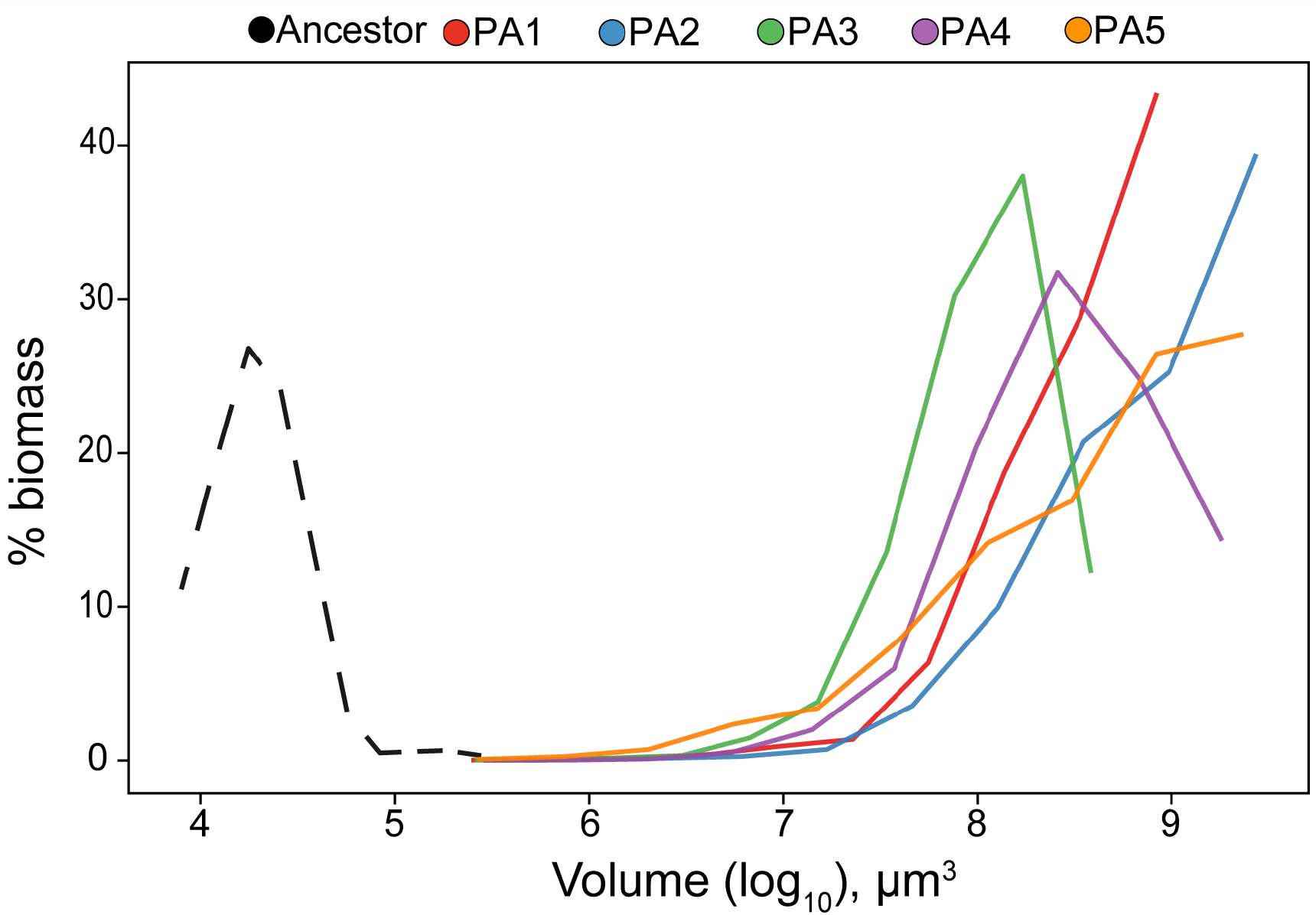
Biomass distribution as a function of group size for the ancestral snowflake yeast (dotted line) and 600 day evolved populations of PA1-PA5. The ‘weighted mean size’ used in Figures 1, 2 and 4 is the mean of the biomass distribution.

**Extended Data Figure 4.**
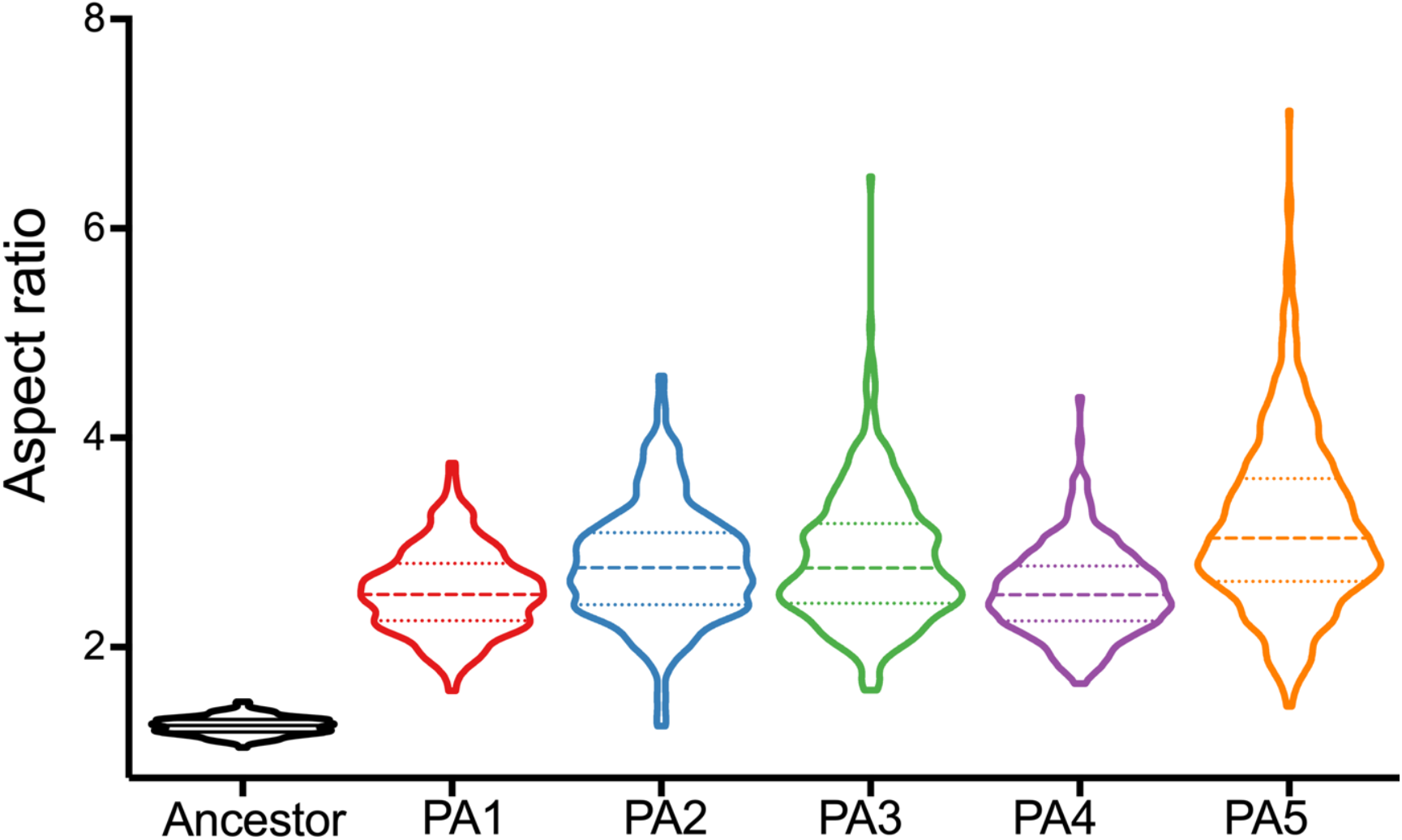
Distribution of aspect ratios for ancestral and 600-day evolved populations of anaerobic snowflake yeast.

**Extended Data Figure 5.**
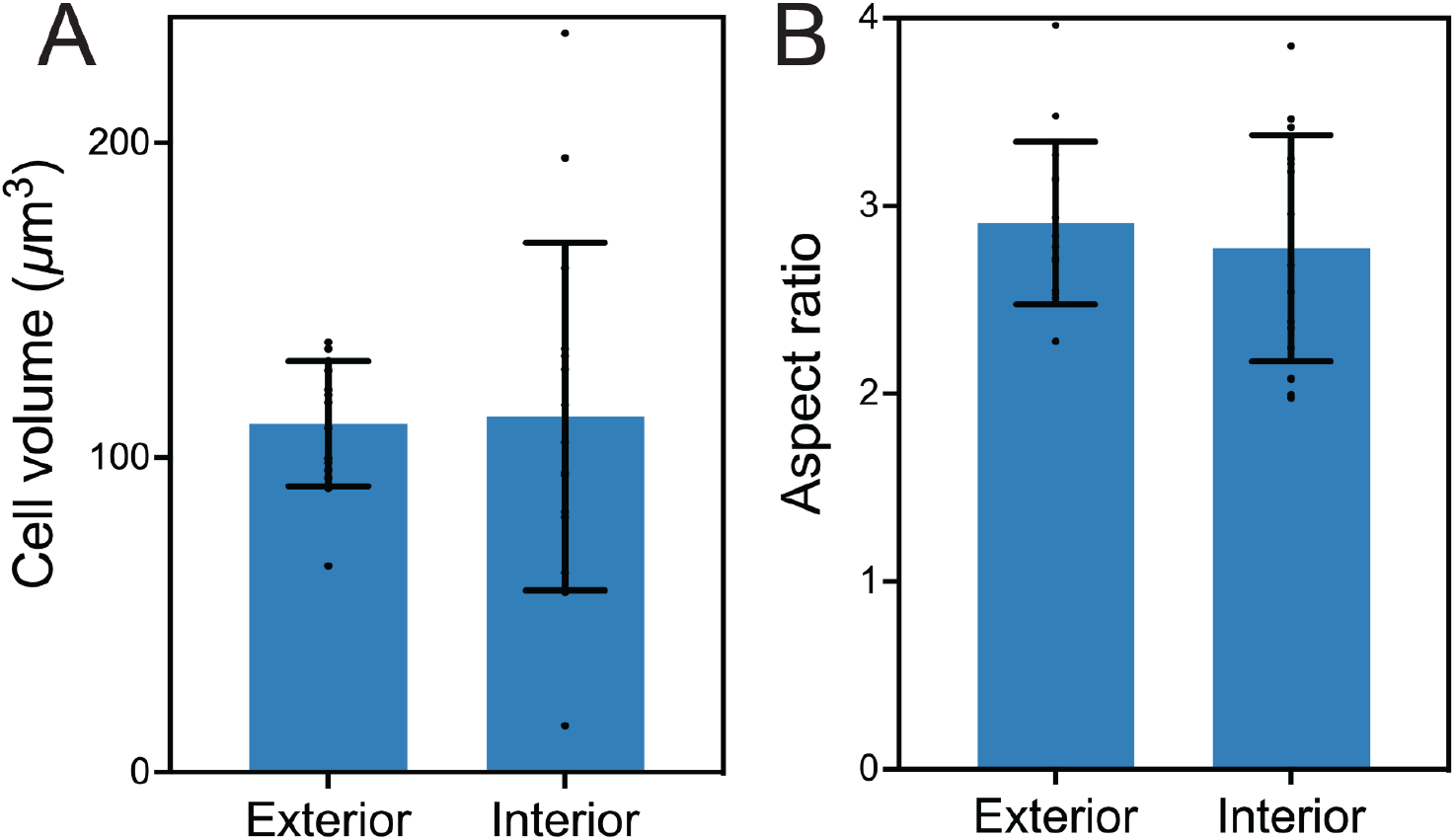
Cell shape is not substantially affected by location within macroscopic yeast. (A) and (B) show cell volume and cell shape (aspect ratio) measured for 10 cells from the interior of a macroscopic cluster and 10 cells from the exterior of a cluster (measured in t600 macroscopic clusters). Average cell volume for exterior and interior are 110.8 μm^3^ and 113.1 μm^3^ (*p*=0.88, *t*=0.15 df=17.55, Welch’s *t*-test), and average cell shape for exterior and interior are 2.9 and 2.8 (*p*=0.51, *t*=0.68, df=14, Welch’s *t*-test). Individual measurements are marked as points, the mean and one standard deviation are indicated by the bar plot.

**Extended Data Figure 6.**
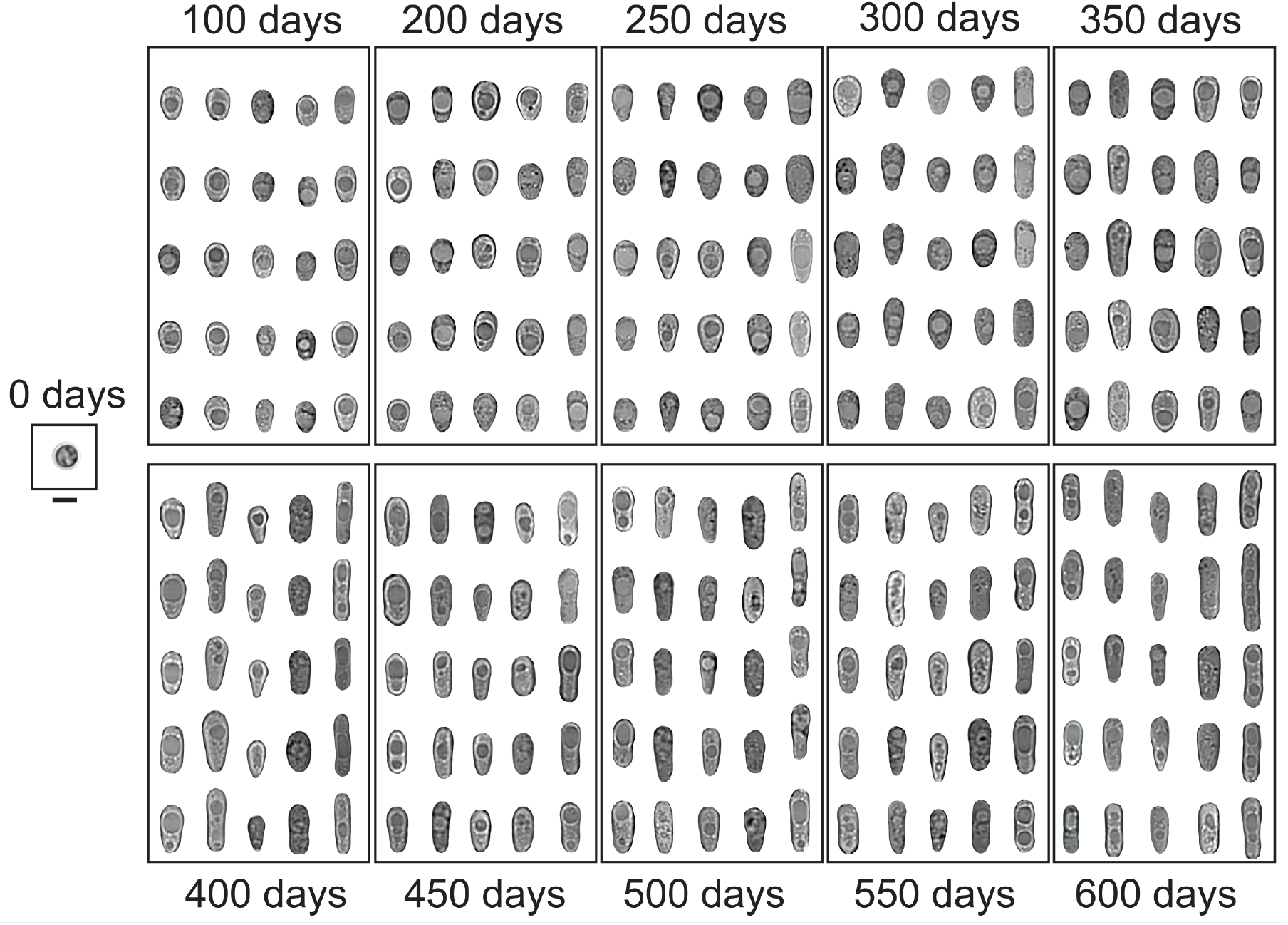
Parallel evolution of elongated cell shape across all five replicates of each PA population. For each evolutionary time point and population, five different cells are shown (organized vertically from left to right: PA1 on the further left and PA5 on the further right in each box). Scale bar is 5 μm (under the ancestral cell). This is a more detailed version of the plot shown in Figure 2c.

**Extended Data Figure 7.**
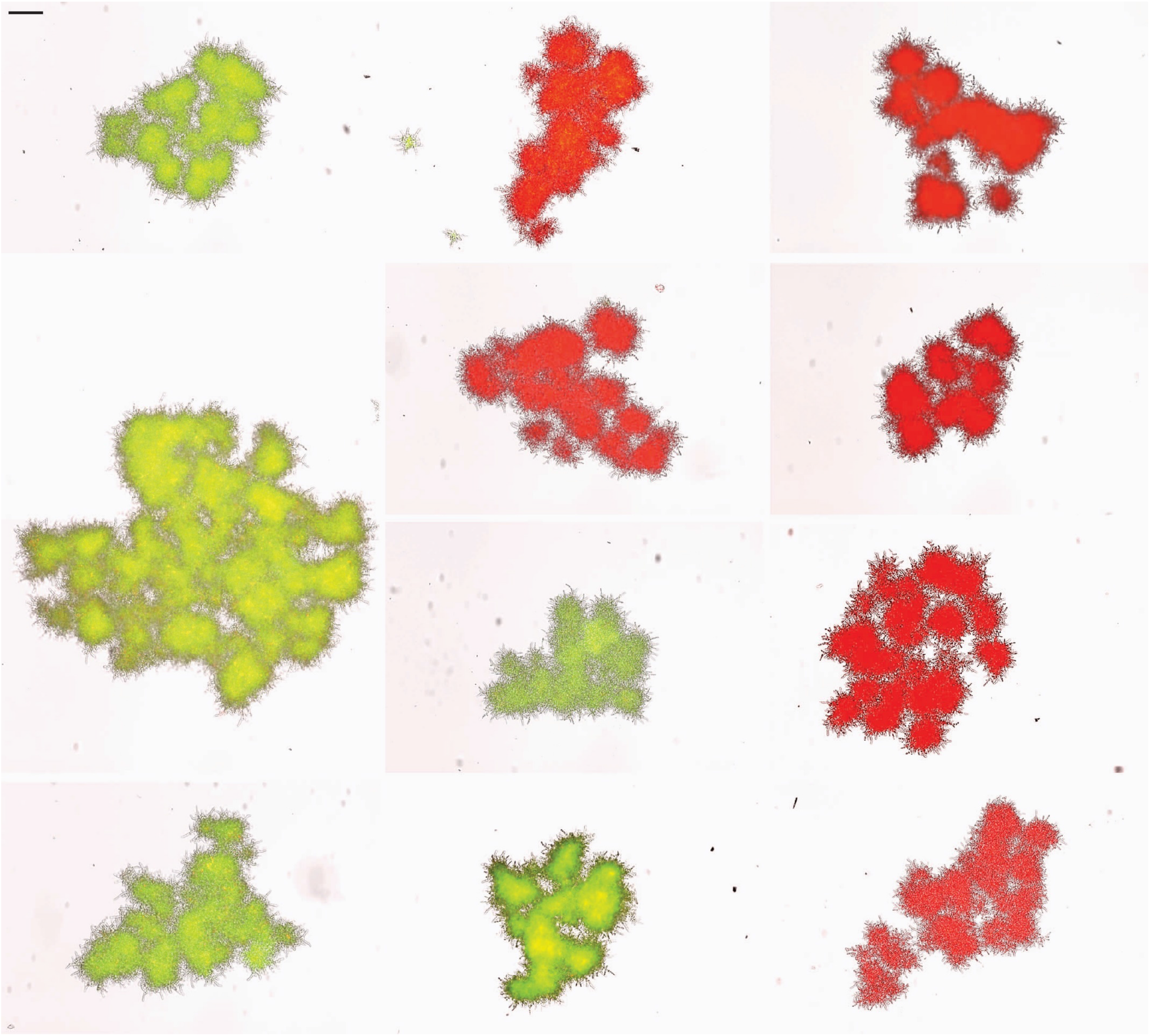
Macroscopic snowflake yeast are monoclonal, growing via permanent mother-daughter cellular bonds, not aggregation. We co-cultured GFP and RFP-tagged genotypes of a macroscopic single strain isolate (PA2, strain ID: GOB1413-600) for 5 days, then imaged 70 clusters on a Nikon Ti-E. Shown are a composite of 11 individual clusters, which all remain entirely green or red. Individual clusters were compressed with a coverslip for imaging, resulting in their fragmentation into multiple modules. Scale bar (top-left) is 100 μm.

**Extended Data Figure 8.**
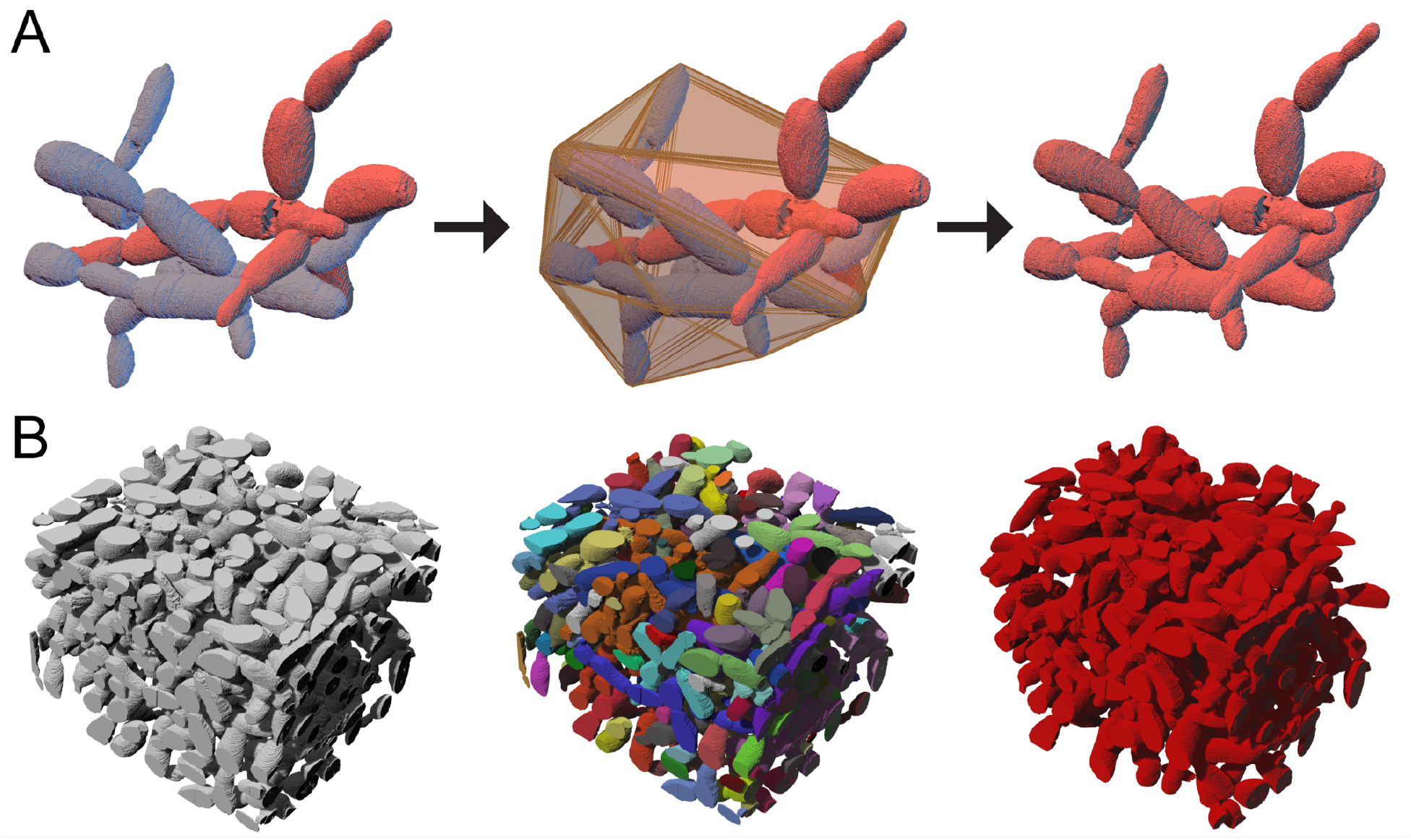
Quantifying entanglement via analysis of the topology and geometry of a snowflake yeast cluster. (A) We measured entanglement of individual components by fitting a convex hull around each component, and determining whether the other component overlaps with the space bounded by this convex hull. Here we just show the convex hull for the blue component, which overlaps with the red component. These components are thus part of the same entangled component. (B) Using this approach, we identified the components within a sub-volume of a macroscopic snowflake yeast, and used a percolation analysis to examine the fraction of the biomass that is part of the same entangled component (colored in red).

**Extended Data Figure 9.**
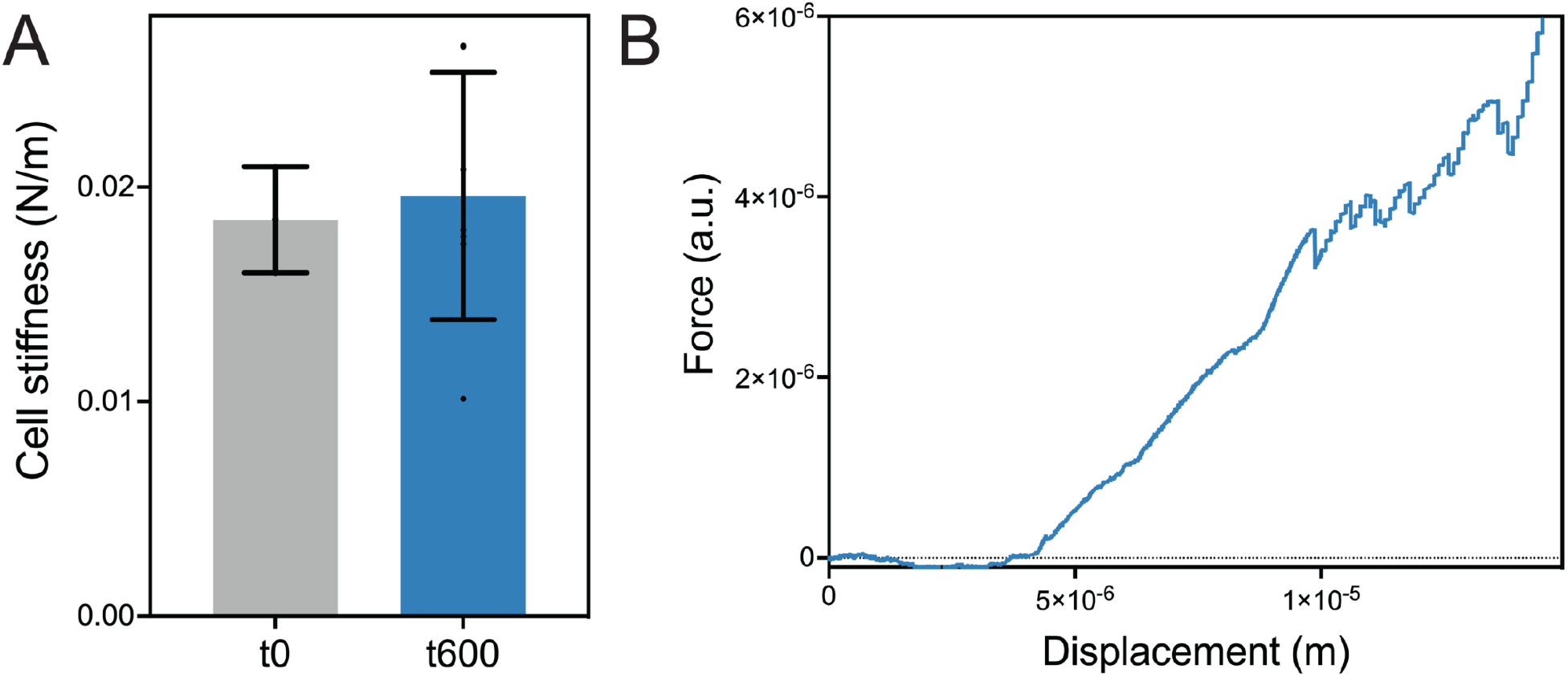
(A) Individual cells do not change their stiffness over 600 rounds of selection (average cell stiffness for the ancestor and t600 isolates are 0.019 and 0.020, respectively. *p*=0.77, t=0.31, df=8, Welch’s unequal variances *t*-test). Single-cell stiffness values measured from atomic force microscopy (AFM) of individual cells. Error bars are one standard deviation. (B) Macroscopic snowflake yeast fractured into small modules prior to compression do not show strain stiffening behavior. Shown here is an AFM trajectory of cantilever deflection vs displacement for one t600 cluster that has been crushed into small, unentangled pieces.

**Extended Data Figure 10.**
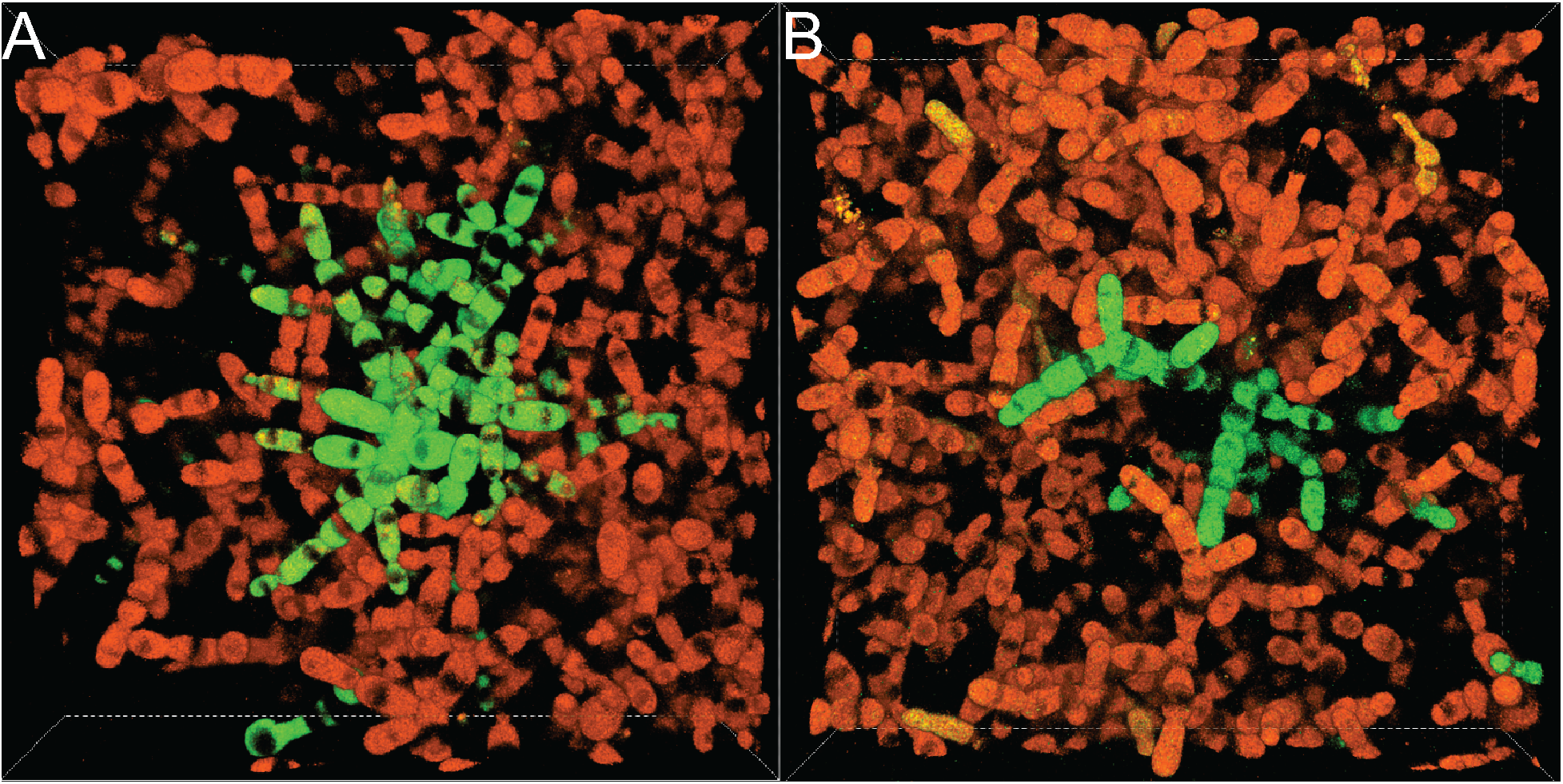
Representative confocal images show chimeric clusters that are formed after growth in liquid culture followed by entanglement on agar plates. Each frame is 139.64 x 139.64 x 34.50 μm in X, Y, and Z axes, respectively.

**Extended Data Figure 11.**
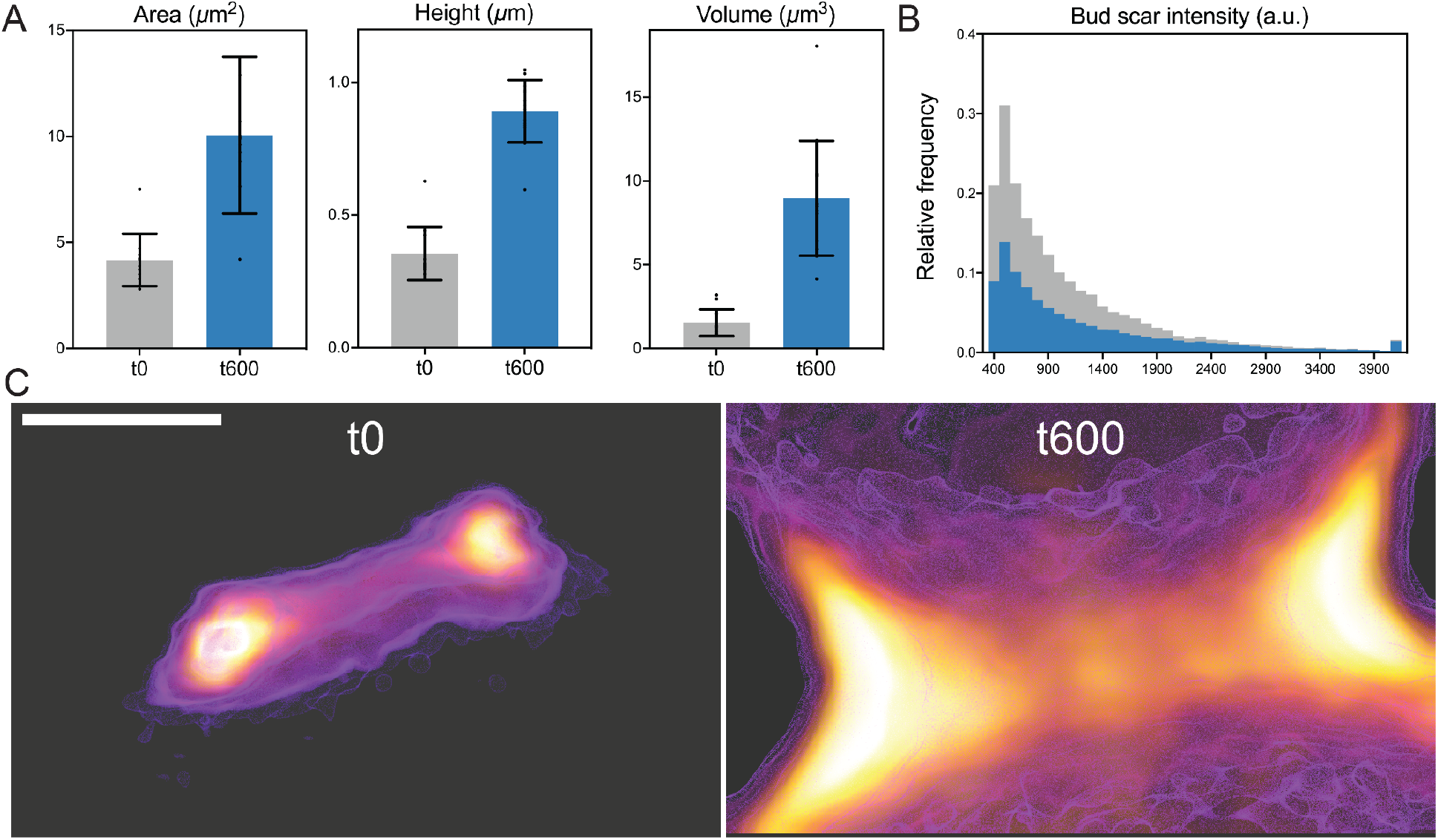
Dimensions of bud scars connecting cells in microscopic, ancestral (t0, gray) and macroscopic, evolved snowflake yeast clusters (PA2 t600, blue). Macroscopic t600 yeast had 2.4x larger bud scar cross-sectional area (A; *t*=5.3, df=24, *p*<0.001), 2.8x greater bud scar height (B; *t*=12.5, df=24, p<0.001), resulting in bud scars with 5.8-fold greater volume (C; *t*=7.3, df=24, *p*<0.001) than the microscopic ancestor. Error bars are one standard deviation. (B) Histogram of pixel intensities for bud scars stained with chitin stain calcofluor white, isolated from ancestor (t0, microscopic) and t600 (macroscopic) bud scars. The t600 strain has a 27% higher mean fluorescence intensity, suggesting that they may have evolved moderately chitin density in the bud scar. (C) The size differences in bud scars is readily visible. Shown are the side view of buds from the ancestor (left) and t600 evolved (right), imaged at the same microscope settings. The scale bar is 0.5 μm.

**Supplementary Movie 1.** Comparison of the ancestor and a population of macroscopic snowflake yeast (PA2-t600, on the right).

**Supplementary Table 1.** A list of primers, plasmids, and strains used in the study.

**Supplementary File 1.** Code for the simple 3D biophysical simulation. These simulations, described at a high level in the methods section of this paper, were adapted from previous work by Jacobeen 2018 and Day 2022 ^31,82^. The code is self-contained and commented. Please reach out to Thomas Day or Peter Yunker with questions.

